# Platelets favor the outgrowth of established metastases

**DOI:** 10.1101/2022.10.28.514195

**Authors:** Maria J. Garcia-Leon, Olivier Lefebvre, Gautier Follain, Clarisse Mouriaux, Ignacio Busnelli, Annabel Larnicol, Florent Colin, Vincent Mittelheisser, Marina Peralta, Catherine Bourdon, Raphael Samaniego, Martine Jandrot-Perrus, Pierre H. Mangin, Jacky G. Goetz

## Abstract

Despite abundant evidence demonstrating that platelets foster metastasis, a therapeutic approach based on anti-platelet agents is not an option due to the risk of hemorrhages. In addition, whether platelets can regulate metastasis at the late stages of the disease remains unknown. In this study, we subjected syngeneic models of metastasis to various thrombocytopenic regimes to show that platelets provide a biphasic contribution to metastasis. While potent intravascular binding of platelets to tumor cells efficiently promotes metastasis, platelets further support the outgrowth of established metastases. Genetic depletion and pharmacological targeting of the platelet-specific receptor GPVI in humanized mouse models efficiently reduced the growth of established metastases, independently of active platelet binding to tumor cells in the bloodstream. Our study is the first to demonstrate therapeutic efficacy when targeting animals carrying growing metastases. It further identifies GPVI as the first molecular target whose inhibition can impair metastasis without inducing collateral hemostatic perturbations.

## INTRODUCTION

Metastasis is a pathological process during which tumor cells (TCs) use the bloodstream to colonize distant organs. One of the most frequent target organs of metastatic cancers are the lungs (1). Around 21–32% of breast cancers are accompanied by lung metastases (2), which dramatically impacts patient mortality (3). Yet, the fundamental mechanisms driving lung metastasis remain poorly understood, limiting the development of efficient therapeutic schemes. When disseminating, circulating TCs (CTCs) face several hostile forces that are detrimental to their survival (4). Yet, they can exploit hemodynamics to arrest and successfully grow upon extravasation (5-7). In addition, CTCs efficiently partner with the bloodstream ecosystem and interact, either as a single or cluster of cells, with blood components on their way to metastasis (8-11).

Among these, platelets are perfect allies. They promote metastatic progression (12) by protecting TCs from destruction by shear forces. They mediate immune evasion, favor adhesion and extravasation at distant sites, facilitate neoangiogenesis and mediate a tumor-prone inflammation (13). Although recent work suggests that platelets can also hamper tumor initiation and growth, as shown in the liver (14-16) and ovarian tumors (14), abundant research thus demonstrates that they mostly promote tumor cell metastasis. A seminal study documented the inhibition of metastasis by platelet depletion, which can be rescued by platelets transfusion (17). The depletion of platelet receptors such as the vWF receptor GPIb, and the collagen receptor GPVI, results in the inhibition of lung metastasis of melanoma and lung carcinoma cells, respectively (18,19).

Anti-platelet strategies became inevitably obvious candidates for impairing metastasis. Yet, they face multiple limitations. First, in the bloodstream, CTCs rapidly bind, activate, and aggregate circulating platelets through the so-called Tumor Cell-Induced Platelet Aggregation (TCIPA). This provides a physical shield to CTCs and favors intravascular arrest and survival (20,21) as well as successful extravasation and metastatic outgrowth (22-24). While it is thus tempting to target the intravascular CTC-platelet interaction with anti-platelet therapies for impairing metastasis, whether this is a universal feature of cancers remains unexplored. It is thus likely that results may differ depending on cancer types (25,26) leading to an unclear outcome when using antiplatelet drugs in oncologic patients (27). It is thus mandatory to explore whether platelets equally bind CTCs depending on the cancer type or stage, so that inhibitory drugs could be developed for standard cancer treatments and/or prophylaxis. Second, our current knowledge of the pro-metastatic function of platelets focuses on their contribution to the intravascular behavior of CTCs, which subsequently impacts metastatic fitness (13,22,23). Whether platelets are capable of tuning other, extravascular and late, metastatic steps remain unknown and unexplored. Yet, they could offer additional means for targeting platelets during metastasis. Finally, the development of efficient anti-platelet strategies to fight metastasis faces major pitfalls that are inherent to the properties of platelets (28,29). The majority the current of anti-platelet strategies are accompanied by increased bleeding risk (30) that prevents using them against metastasis. This remains a major concern and explains why anti-platelet therapies could not be used in the clinic.

Building on such limitations, we designed a study providing a comprehensive characterization of the interaction efficiency between platelets and a panel of metastatic cells. Selected TCs with different platelet-interacting abilities were subjected to several thrombocytopenic (TCP) regimes in mouse models to interrogate the timings at which platelets mostly contribute to metastatic outgrowth. We show that platelets binding efficiencies are crucial for the adhesion and survival of TCs within the vasculature and we provide evidence that platelet binding profiles scale with their long-term metastatic potential *in vivo*. We observed that platelets efficiently colonize metastatic foci, in both mouse and human samples. Depletion of platelets in animals carrying growing metastases was sufficient to reduce metastatic outgrowth demonstrating, for the first time, that platelets favor metastasis independently of platelet-tumor cell interaction in the bloodstream, which could be targeted with new treatment strategies. We focused our attention on the platelet receptor GPVI, which is exclusively expressed at the surface of platelets and whose targeting inhibits platelet function with no impact on bleeding time. Genetic depletion and pharmacological targeting of the platelet-specific receptor GPVI in humanized mouse models efficiently reduced the growth of established metastases, independently of active platelet binding to TCs in the bloodstream. Our findings highlight, for the first time, that interfering with GPVI can impair metastatic outgrowth of already established metastases. We believe that such discovery paves the way for establishing successful therapeutic intervention in patients where the primary tumor has already generated one or several metastatic sites.

## RESULTS

### Tumor cells bind and activate platelets *in vitro* with different efficiencies

Platelets have been reported to bind circulating TCs (12) but whether this applies to every cell type remains unknown. We first checked *in vitro* the efficiency of human platelets to interact with various human and mouse TC lines, either from breast cancer (epithelium-derived) or melanoma, using scanning electron microscopy (SEM). We found out that platelets do not equally bind TCs (Fig.1A-B). While some cells such as 4T1 or MCF7 efficiently bind platelets, others such as B16F10 and MDA-MB-231 cannot bind more platelets than passive beads (Fig. 1A-B, S1A). We assessed the morphology of bound platelets (Fig.1C-D) and observed that TCs behave as weak platelet agonists. We probed their activation through their ability to promote TCIPA (Fig.1E-F) which correlated with platelet binding. We further observed that TCs efficiently bind platelets in the absence of plasma proteins (Fig.S1E-F) but fail to promote aggregation (Fig.S1B,D) or activate them (Fig.S1G-H), altogether indicating that TCs do not equally bind and activate platelets, raising the possibility that both processes directly impact the metastatic properties of TCs. To test such a hypothesis, we selected two mouse TC lines with opposite binding and platelet activation behavior (4T1 vs B16) that are compatible with experimental metastasis approaches in a syngeneic background.

**Fig. 1.**
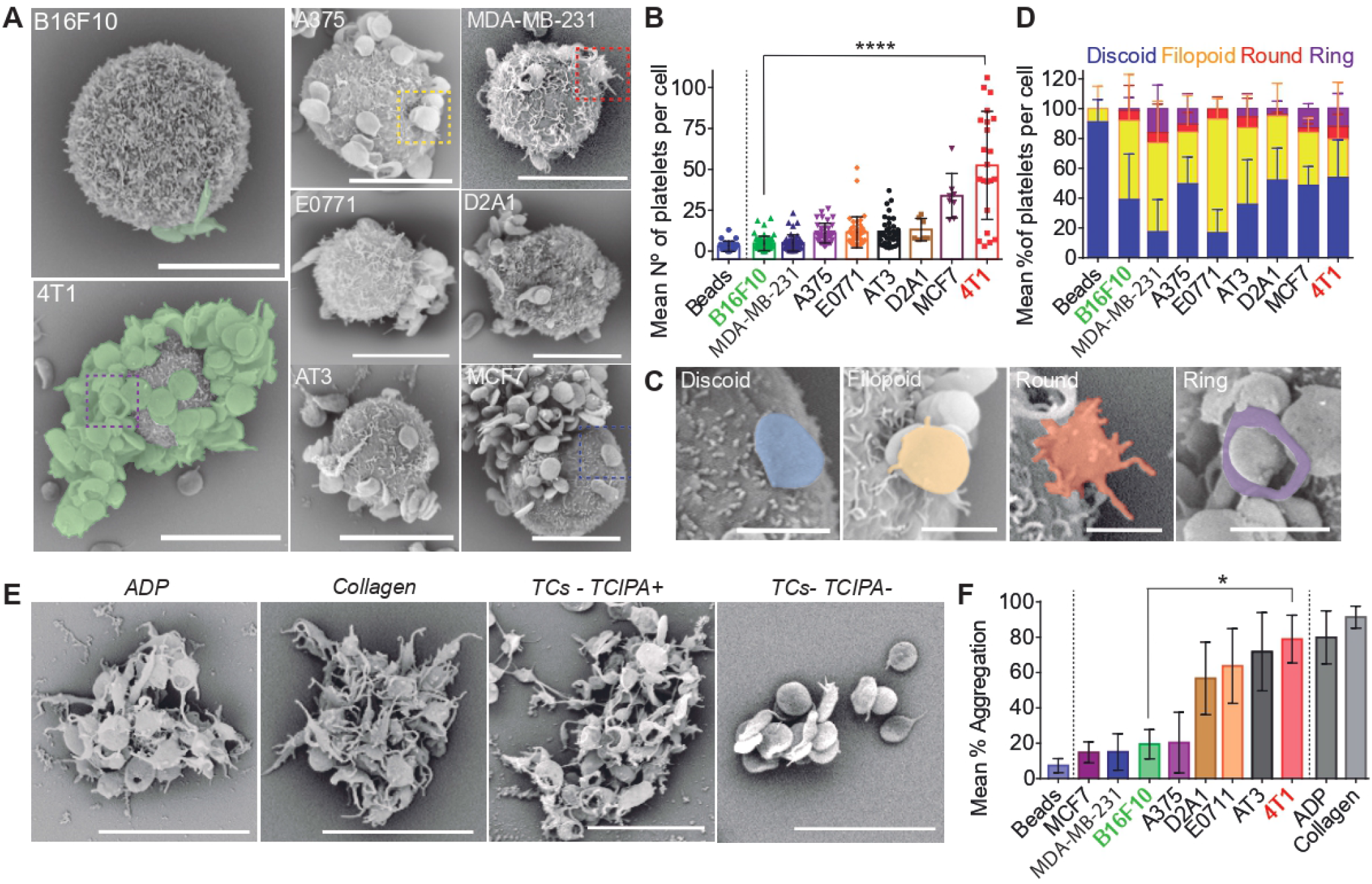
TCs bind and aggregate blood platelets with different efficiencies. (A) SEM images of TCs interacting with mouse platelets contained within mouse citrated platelet rich plasma (cPRP). Mouse breast carcinoma (4T1) and mouse melanoma (B16F10) cells are highlighted, as they show the most different platelet binding profiles (high versus low, respectively). (B) Platelet binding profiles quantification graph of (A). Number of platelets per cell as mean +/-SD from 2 to 5 independent experiments. (C) SEM images of TC-bound platelets at different activation states including: discoid (resting), filopodia (low activation) and round (full activation). Ring-shaped platelets of unknown function were also observed in a minor frequency. (D) Mean relative number (%) of each platelet shape per TC; activation stages are color-coded. (E) SEM images of mouse platelet aggregates obtained after the 30 minutes of platelet aggregation assay using cPRP with ADP and collagen as classic agonists, and 4T1 and B16F10 TCs TCIPA+ or TCIPA-cells. (F) Relative (%) mean human cPRP platelet aggregation in tested TCs. Representative data from a total of 7 to 50 cells per cell line from 2 to 4 independent experiments showed as mean +/-SD.

### Platelets orchestrate the intravascular behavior of circulating TCs and control lung seeding *in vivo*

It is well-established that the early seeding of TCs shapes their metastatic potential (31) so we next interrogated whether differential platelet binding may influence metastasis *in vivo*. We designed an experimental lung metastasis assay that allows us to probe the behavior of TCs at very early and sequential steps of the metastatic cascade (Fig.2A). We depleted platelets using an α-GPIb treatment that proved to be equivalently efficient in both animal models (Balb/c and C57BL/6, Fig.2B) and assessed TCs lung seeding by sequential *in vivo* imaging of 4T1 or B16F10 luciferase-expressing cells, starting from 15 minutes post-injection (mpi) up to 24 hours post-injection (hpi) (Fig.2A). High-bind 4T1 cells reached and efficiently seeded control lungs upon clearance from 15 mpi to 24 hpi (33-fold), as shown previously for metastatic cells (31). However, when 4T1 cells were injected in thrombocytopenic mice, lung seeding was drastically reduced by 10-fold when compared to control conditions. In contrast, the lung seeding potential of low-bind B16F10 cells was low when compared to 4T1 and independent of the presence of platelets (Fig.2C-D). These data suggest that lung seeding scales with the platelet binding potential of TCs.

**Fig. 2.**
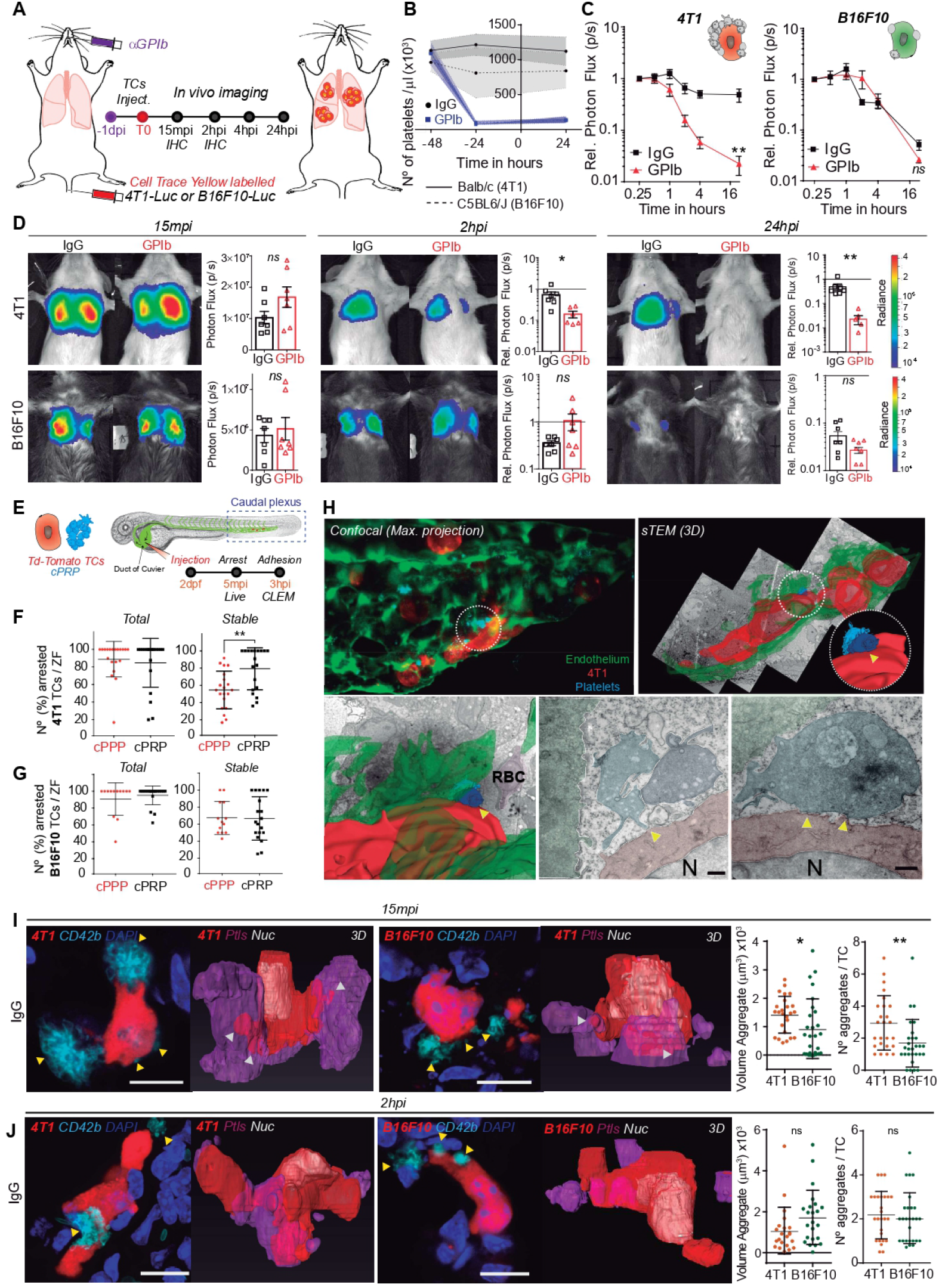
*In vivo* lung seeding of 4T1 or B16F10 TCs is differentially influenced by platelets. (A) Infographics showing the experimental setting to study the early steps (24h) of the metastatic cascade in mice in the presence or absence of platelets. Cell Trace-labelled 4T1-Luc or B16F10-Luc TCs are injected intravenously in normal or thrombocytopenic (TCP) mice (IgG- or GPIb-treated, respectively) 1 day post-treatment (1dpi). Lung seeding is followed by *in vivo* bioluminescence and analysis of arrested single cells is performed by immunohistochemistry at 15 minutes port-injection (15mpi) and 2 hours post-injection (2hpi). (B) Graph showing platelet counts on IgG- and GPIb-treated Balb/c and C5BL6/J animals. (C) Kinetics of relative (to input) bioluminescent 4T1-Luc and B16F10-Luc TCs lung seeding during 24 hours in normal and TCP mice. Data represented as mean +/-SEM, 2 independent experiments of n=3 to 4. (D) Representative images and time point comparisons of relative lung bioluminescence in control versus TCP animals in 4T1 (up) and B16F10 (down) TCs models. Data are represented as mean +/- SEM, Mann-Whitney analysis. (E) Infographics showing the experimental setting to study the early steps of the metastatic cascade *in vivo* in the ZF embryo. Td-tomato-expressing 4T1 or B16F10 cells are intravenously microinjected into the ZF embryo via the duct of Cuvier, together with human cPPP (control) or human cPRP. TCs circulation and arrest are imaged immediately by live video microscopy. Stable adhesion and TC-platelet interactions are imaged by confocal microscopy at 3hpi. (F) Quantification of total and stable arrest of 4T1 cells in the caudal plexus of the ZF embryo during 5 minutes of live video microscopy. Data represented as mean +/- SD, 4 independent experiments with a minimum of 15 ZF embryos per experiment. (G) Quantification as in (F) for B16F10 cells. (H) Representative confocal and CLEM images of 4T1 cells arrested intravascularly in the caudal plexus of the ZF embryo after 3hpi. Arrowheads indicate contact between platelets and TCs. Asterisks indicate contact between platelets and endothelium. Scale: 500nm. 5 to 10 ZF were injected with 4T1 cells in 3 independent experiments; only one representative fish was selected for CLEM analysis. (I) Representative confocal images of single 4T1 or B16F10 cells arrested in control and TCP mouse lungs at 15mpi. Arrowheads show the interaction of single platelets and platelet aggregates with the TCs. Scale bar: 10µm. On the right, the volume and number of platelet aggregates around the arrested TCs is shown. Platelet and TCs volumes were calculated by TC and platelets segmentation in AMIRA. Data represented as mean +/- SD, 1 independent experiment with n=3 different mice per group and no less than 10 single-cell images per mice. (J) Representative confocal images of single 4T1 or B16F10 cells arrested in normal and TCP mouse lungs at 2hpi. Arrowheads show the interaction of single platelets and platelet aggregates with the TCs. Scale bars 10µm. On the right, the volume and number of platelet aggregates around the arrested TCs in normal and TCP mice. Same analysis as in (I).

To provide a live and quantitative description of this platelet-dependent intravascular arrest of TCs, we next used our previously-established experimental metastasis model in the zebrafish embryo (5). Fluorescently-labeled TCs and human citrated platelet-rich plasma (cPRP) were co-injected in the duct of Cuvier of *Tg(Fli1a:GFP)* embryos at 2 days post-fertilization (dpf) and live-monitored during 4 minutes post-injection (mpi) (Fig.2E). Careful live tracking of TCs5, showed that platelets favor the stable arrest of 4T1 in the caudal plexus (Fig.2F, S2A) thus reducing their circulation time (Fig.S2A). In contrast, platelets did not affect the circulation and arrest patterns of B16F10 (Fig.2G, Fig.S2B). A heat mapping analysis that allows quantitatively locating the hotspot of the arrest of TCs, revealed that platelets favored the stable arrest of 4T1 cells in high shear arterial vessels (DA, dorsal aorta; CV, caudal vein) without impacting the entry of TCs into small sized vessels (ISV, inter-somitic vessels, Fig.S2A,C,D) where TCs mostly arrest through physical constraint imposed by blood vessels. Interestingly, although they did not affect the number of stable arrest events occurring for B16F10 in non-constraining arterial vessels, platelets favored their entry and arrest into inter-somitic vessels (Fig.S2B,C,D). We next probed arrest events at the nanoscale using correlative light electron microscopy (CLEM) (5,32) and observed that stably arrested 4T1 cells efficiently interacted with platelets with an activated morphology (filopodia, Fig.2H) and endothelial cells. Altogether, this provides the first *in vivo* demonstration that platelet binding favors the intravascular arrest and adhesion of circulating TCs to the endothelial wall, which can be probed at nanoscale (CLEM).

To further document *in vivo* this unequal binding (Fig.1), we analyzed TC-platelet binding in mouse lung sections containing early-seeded (15 mpi and 2 hpi) 4T1 or B16F10 TCs (Fig.S3A). While imaging at 15 mpi allows catching the very initial *in vivo* behavior of TCs, around 50% of the input cells are already cleared from the lungs after 2 hpi in the absence of platelets (Fig.2C-D), thus identifying a critical platelet-dependent step. 3D high-resolution imaging at 15 mpi revealed that 4T1 cells bind, recruit and aggregate more platelets than B16F10 also *in vivo* (Fig.2I), validating our initial *in vitro* observations. As expected, the size (volume) and number of platelet aggregates were also significantly lower in thrombocytopenic animals (Fig.S3B). At 2 hpi, the number of platelets found near B16F10 cells increased significantly (Fig.2J) suggesting that platelets are passively recruited *in vivo* to arrest sites, by either blood flow clogging, TC cytoplasmic release of platelet aggregation inducers, or other mechanisms. When probing cell survival using cell morphometrics at 2 hpi, we observed increased nuclear fragmentation (non-viable cells) in 4T1 cells of α-GPIb-treated animals, as well as in low-bind B16F10, independently of the presence of platelets. (Fig.S3C-D). This effect was not due to distinct survival behaviors between the two cell lines, as both displayed equivalent *in vitro* viability up to 90 min (Fig.S3A, bottom), but rather linked to the presence of platelets intravascularly (Fig.S3D, bottom) suggesting that platelets favor intravascular survival of 4T1 cells, as previously reported (20). Interestingly, B16F10 cells, that do not actively bind platelets, *in vitro* and *in vivo*, can passively recruit platelets at 2 hpi (Fig.S3E) suggesting that they could favor the intravascular survival of neighboring viable arrested cells, evidence in line with previous observations (33).

In conclusion, using a novel *in vivo* analysis of early lung metastatic seeding in mice and zebrafish embryos, we show that platelets favor early seeding (15 hpi) and survival of TCs able to actively bind to them (4T1 cells). Additionally, the arrest and intravascular death of low platelet binding capacity TCs (B16) allow the passive and later recruitment of platelets (> 2hpi) that can support survival of neighboring viable cells. As such, however, platelets do not equally promote metastatic seeding as 4T1 cells seed the lungs more efficiently than B16F10 cells (Fig.2D).

### Platelets control metastatic fitness at early stages

Having demonstrated that platelet binding efficiency correlates with intravascular arrest and survival driving efficient lung seeding, we next wondered whether such early steps would translate into efficient metastatic outgrowth. We longitudinally tracked metastatic burden over time in control and thrombocytopenic mice after a single treatment with α-GPIb antibody prior TCs injection (14 days, Fig.3A). Because platelet counts quickly recovered and reached approximately 50% of the normal numbers at day 7 (Fig.3B), we name this model as short-term thrombocytopenia (shortTCP). Metastatic outgrowth (day 14) was significantly reduced by shortTCP in both models (Fig.3C). Interestingly, while the overall effect of shortTCP on 4T1 metastatic outgrowth was a consequence of its effect on early lung seeding (24 hpi and Fig.2), shortTCP impacted the growth of B16F10 metastases only after lung seeding. Indeed, although B16F10 lung seeding was equivalent at 24hpi (Fig.2C-D, 3C-D), the metastatic burden was significantly reduced by shortTCP (14dpi, Fig.3C-D). While we cannot exclude that metastatic outgrowth is impacted by the quick recovery of the platelet counts (Fig.3B), these data demonstrate that platelets control early events of metastatic outgrowth of 4T1 (day 1 post-injection) that later translate into a significant reduction of metastatic burden (day 14). Yet, these data also hint at an additional effect of platelets as can be observed in the low-bind B16F10 model where thrombocytopenia has no effect on lung seeding (day 1) and yet translates into significant metastasis impairment at day 14.

**Fig. 3.**
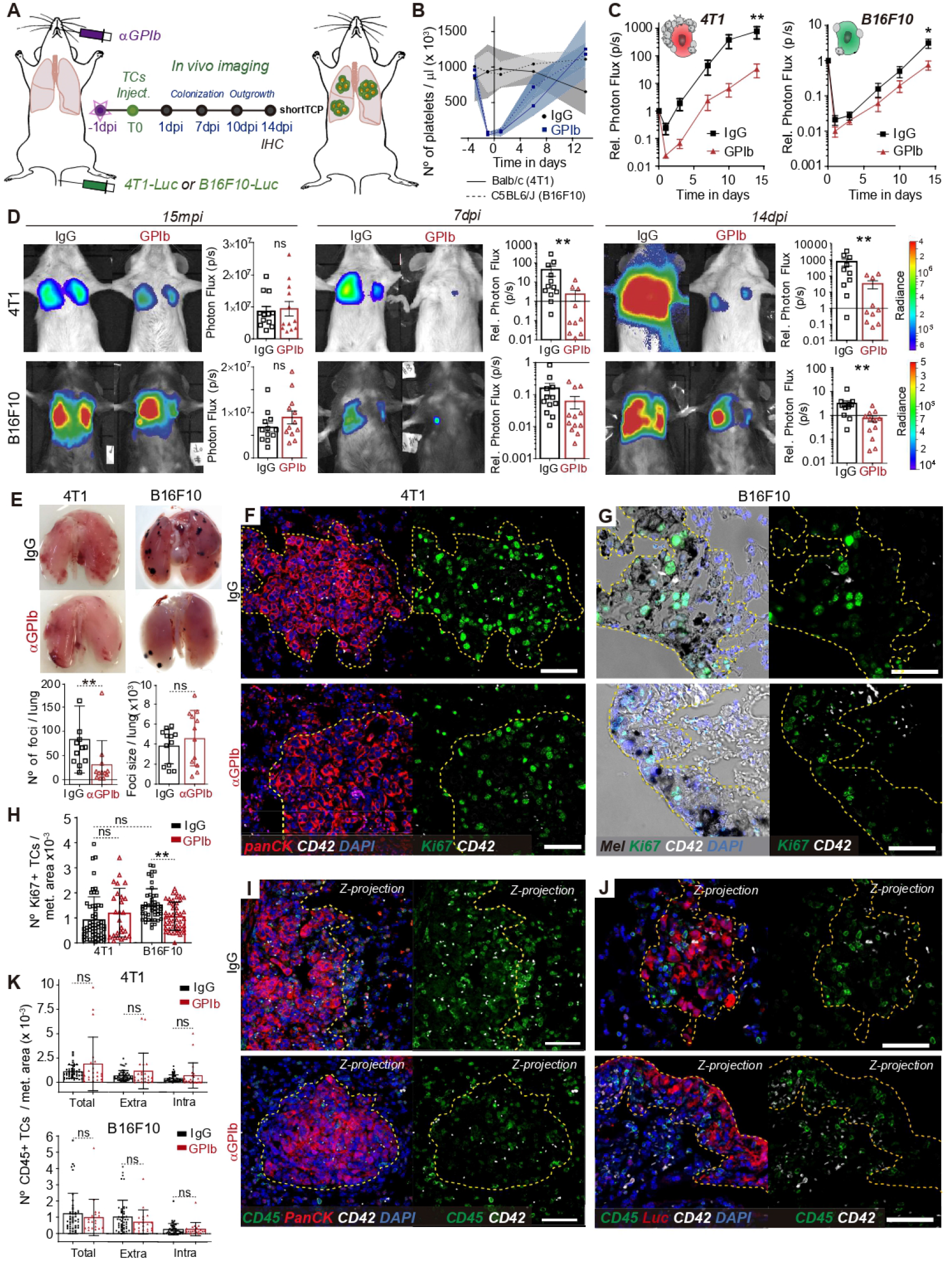
Long-term TCs metastatic potential is defined by the early TC–Platelet interplay. (A) Representative images and time point comparisons of total lung bioluminescence in normal versus TCP mice in the 4T1 (up) and B16F10 (down) TCs models used for IF analyses of TC intravascular arrest. Two cohorts of mice were used, the first was sacrificed at 15mpi (upper panel) and the second at 2hpi (lower panel). Data are represented as mean +/- SD of 6 mice from 1 experiment. Bottom left, relative *in vitro* viabilities of 4T1 and B16F10 cells up to 90min. Bottom right, live and fluorescence imaging of and PFA-fixed / OCT-embedded 4T1 TCs labelled with CellTrace Yellow and DAPI. (B) Representative AMIRA segmentation images of single TCs arrested intravascularly in the mouse lungs after 15 minutes post-injection (upper panel) and 2hpi (lower panel). Scale: 10µm. Violin plots represent the number of aggregates per TC and the volume of aggregates per TC calculated using AMIRA segmentation of confocal images. Data are shown as mean +/- SD. (C) Violin plots depicting TCs volumes calculated in AMIRA to obtain an estimation of cell viability. Same analysis as in (B). (D) Representative images of lung-arrested non-viable B16F10 cells, as shown by the absence or fragmented nucleus. Scale: 10µm. Bar graphs (bottom) show the frequency (%) of viable B16F10 cells at 15mpi and 2hpi calculated by nuclear integrity. One experiment with n=3 different mice per group and no less than 10 single-cell images per mice. Data are shown as mean +/- SD. (E) Right: Representative confocal images of intravascular B16F10 clusters at 2hpi depicting viable cells co-opted together with non-viable cells and large platelet aggregates. Scale: 10µm. Left: Violin plots showing the volume of platelet aggregates around viable (V) and non-viable (NV) B16F10 at 15 mpi and 2phi. Same analysis as in (B).

To explore the mechanisms that may be operating after seeding, metastatic lungs were surgically resected on day 14. Macroscopic inspection of B16F10 metastatic foci revealed, as expected, that shortTCP significantly reduced their numbers (Fig.3E, Fig.S4C), in agreement with the lower bioluminescent signal observed at day 14 (Fig.3C-D). While the size of the B16F10 foci appeared similar (Fig.3E), these lesions displayed a significant decrease in Ki67-positive cells indicating that their proliferation index was impacted by platelet depletion (Ki67, Fig.3G,H). Such effect was not observed in 4T1 foci (Ki67, Fig.3F,H, Fig.S4C) suggesting that platelets may tune proliferation of B16F10 during the growth of already-established metastatic foci. While platelets were shown to favor the recruitment of a pro-inflammatory microenvironment favoring the proliferation of TCs (34), short-TCP did not affect immune cell recruitment in any of the two cell models (Fig.3I,J,K).

Platelets were shown to colonize and support primary tumors (14,34,35). Yet, whether they also populate and support metastatic foci remains to be demonstrated. We assessed the number of single (< or = 3 platelets) or aggregated (>3 platelets) intrametastatic platelets and observed that total platelet numbers and aggregates were reduced upon shortTCP in both models (Fig.S4A), despite the recovery in platelet counts (Fig.3B). Importantly, while platelet numbers within control 4T1 and B16F10 metastases were equivalent (Fig.S4B), the number of aggregates (>3 platelets) within B16F10 metastases was significantly higher than in 4T1 metastases, suggesting that B16F10 cells are likely to better recruit, activate and aggregate platelets within metastatic foci (Fig.S4B). Altogether, these results demonstrate that transient thrombocytopenia perturbs early seeding and metastatic outgrowth of TC that strongly binds platelets, while it does not effect on the seeding of TC that poorly bind platelets intravascularly, yet it perturbs their subsequent outgrowth. Because platelet aggregates populate metastatic foci and shape their proliferation index, it is tempting to speculate that they can shape late stages of metastasis.

### Platelets control late steps of metastatic outgrowth

Because platelets populate metastatic foci (Fig.S4A), release important pro-metastatic molecules (36,37) and to circumvent the limitations of the yet classically used shortTCP model, we designed a long-term thrombocytopenia model (longTCP) by the sequential injection of α-GPIb mAb every three to four days, starting 24h before TCs injection (Fig.4A). Doing so, we could maintain low platelet counts up to day 10 post-injection (Fig.4B) and therefore fully impair platelets interactions during the initial growth phase of micro-metastatic foci. Metastatic outgrowth of 4T1 cells was similarly reduced in longTCP (Fig.4C-D) when compared to shortTCP (Fig.3C-D) confirming that platelets strongly contribute to the early steps of the metastasis cascade in that model. In sharp contrast, longTCP massively impaired metastatic outgrowth of B16F10 after 24hpi (Fig.4C-D). While shortTCP reduced metastasis by 3-fold, longTCP reduced it by 45-fold, further emphasizing the additive effect of depleting platelets at late stages of metastasis. While lungs were equally seeded in the presence and absence of platelets (Fig.4C,D, 15 mpi), as expected from previous experiments, longTCP completely abrogated the growth of B16F10 cells resulting in metastases-free lungs at 14dpi (Fig.4C, D, E, Fig.S5B). When combined with the observation that platelets are passively recruited by B16F10 cells at 2 hpi (Fig.2) and that they massively populate metastatic foci at 14dpi (Fig.3), this result suggests that platelets may tune metastatic outgrowth independently of the early intravascular interaction with TCs.

**Fig. 4.**
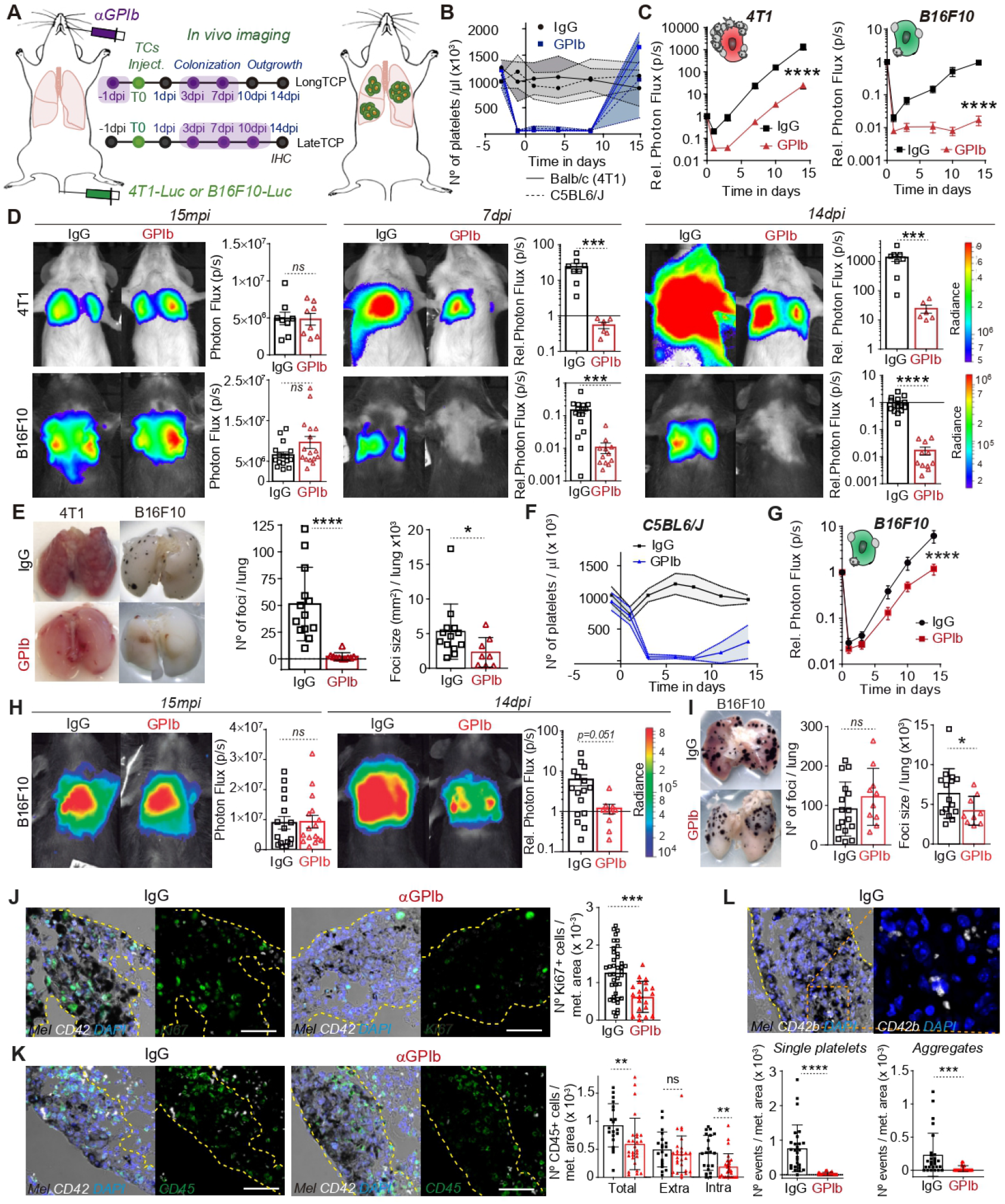
Platelets significantly influence the late outgrowth of B16F10 metastatic foci but not 4T1 ones. (A) Infographics showing the experimental setting to study the late events of the metastatic cascade (14 days) in the presence or absence of platelets. 4T1-Luc or B16F10-Luc TCs are injected intravenously in normal or thrombocytopenic (TCP) mice treated with 3 injections of IgG isotype control or the platelet-depleting antibody αGPIb at different time points. (B) A first set of mice were treated before and after 4T1 or B16F10 TCs to maintain platelet counts low during the whole experiment (longTCP). (C) Bioluminescence kinetics of 4T1-Luc (left) and B16F10-Luc (right) TCs lung seeding and outgrowth in control and long-term TCP mice. Data are presented as mean +/- SEM, 2 independent experiments of n=8. (D) Representative images and time point quantifications (15mpi, 7dpi and 14dpi) of total lung bioluminescence in control versus longTCP animals in 4T1 (up) and B16F10 (down) TCs models. Data are presented as mean +/- SEM. (E) Macroscopic analysis of mice lungs at 14dpi. Pictures of normal and α-GPIb-treated lungs with 4T1 or B16F10 metastatic foci are shown in the left panel. Graphs show B16F10 foci number (middle) and mean foci size (right). Mean +/- SD of 8 mice per group from 1 (4T1) or 2 (B16F10) independent experiments. (F) Following (A), a second set of mice were treated after B16F10 TCs injection, to deplete platelets after TCs seeing in the lungs (lateTCP). (G) Kinetics of B16F10-Luc TCs lungs outgrowth in control and lateTCP mice. (H) Representative images and time point comparisons (15mpi and 14dpi) of total lung bioluminescence in control versus lateTCP animals in B16F10 TCs models. Data are presented as mean +/- SEM, 2 independent experiments of n=8. (I) Macroscopic analysis of mice lungs at 14dpi. Images show normal and α-GPIb-treated lungs with B16F10 metastatic foci. Graphs show B16F10 foci number (middle) and mean foci size (right). Data are presented as mean +/- SD, 8 mice per group from 1 experiment. (J) Immunohistochemical analysis of control and lateTCP mice lungs at 14dpi seeded with B16F10 TCs. Panel shows representative images of proliferative (α-Ki67, green, left) and graph quantification of the number of KI67+ TCs (right) from 8 to 10 images per mice from 2 to 3 mice, 2 independent experiments.(K) Representative images of CD45 immune positive cells (α-CD45, green, left) and graph quantification (right) of CD45+ cell numbers presented as total, extrametastatic (surrounding) or intrametastatic CD45+ cells, from 8 images per mice from 2 to 3 mice, 2 independent experiments. Data are presented as mean +/-SD. (L) Representative images of platelets inside lungs metastases (α-CD42b, white, top) and graph quantification of platelet numbers per metastatic area (bottom) as single platelet (< or = 3) or aggregates platelets (>3) from 8 to 10 images per mice from 2 to 3 mice, 2 independent experiments, mean +/-SD.

To further test this hypothesis, we depleted platelets after the establishment of B16F10 metastatic foci. We decreased platelet counts from 3dpi with three successive injections of α-GPIb until 10dpi (Fig.4A, bottom), naming this novel approach as lateTCP. By controlling platelet counts after the initial growth of metastatic foci, we interrogated a novel contribution of platelets to metastasis (Fig.4F). LateTCP significantly reduced B16F10 metastatic outgrowth from day 7 (Fig. 4G-H), thus confirming that can platelets control the growth of already-established and growing metastatic foci. While the number of foci in control and α-GPIb-treated mice was similar, their size was significantly reduced (Fig.4I, Fig.S5C) as expected from the effect of lateTCP on B16F10 cell proliferation index (Fig.4J,M). As expected, we again observed that the number of intrametastatic platelets was significantly reduced in lateTCP (Fig.4L) suggesting that recruitment and density of platelets within metastatic foci are crucial for their outgrowth. Interestingly, the number of intrametastatic, but not extrametastatic, CD45-positive immune cell infiltrates was significantly reduced in thrombocytopenic animals, suggesting that platelets may modify the immune tumor microenvironment towards immunosuppression (Fig.4K), as previously shown (38). To further support the pro-metastatic contribution of platelets to established metastatic foci, we next interrogated whether platelets would also populate human metastatic foci. With that aim, we analyzed human lung metastasis biopsies from metastatic melanoma patients and found that platelets could easily be detected (Fig.S5A), providing the first evidence that platelets can populate human metastatic foci and further suggesting that they may control metastasis at the late stages of their progression.

### Platelets control late steps of metastasis via GPVI

The late contribution of platelets to metastatic outgrowth opens a promising therapeutic window that could circumvent the current limitations in impairing metastatic extravasation or in detecting early metastatic foci that often occurs even before cancer detection (39,40). Yet, one needs to identify the molecular targets involved and design a strategy that would avoid deleterious thrombocytopenic effects while impairing the outgrowth of established metastases. To do so, we first exploited in-house genetic models that allow us to test the contribution of a specific platelet receptor, namely the collagen and fibrinogen receptor GPVI. GPVI is an immunoglobulin (Ig)–like glycoprotein which binds to many different adhesive proteins and that (41) stimulates platelet activation via GPIIb/IIIa integrins. Recent studies placed it as an attractive candidate in the promotion of platelet-dependent metastasis (42,43), notably because of the minimal impact of its depletion on bleeding. Whether it also tunes late steps of metastasis remains to be demonstrated.

To allow for syngeneicity in the GPVI-/- mouse model (Fig.5A), we engineered the mouse breast cancer AT3 (44) cell line for stable luciferase expression to mirror the previously used 4T1 model, as these cells also displayed a high platelet-binding capacity (Fig.1A). Interestingly, the metastatic outgrowth of high-bind AT3 cells remained unaltered in the absence of GPVI (Fig.5B-C). Similarly, when targeting GPVI pharmacologically with a GPVI-depleting antibody (Fig.S6A,D) that allows for maintaining normal platelet counts (Fig.S6A,B), lung seeding (1dpi) and metastatic outgrowth (14dpi) of high-bind 4T1 cells remained unaffected (Fig.S6C,E-F) suggesting that targeting GPVI might be inefficient for TC that strongly bind platelets and whose metastatic behaviour is shaped by early intravascular platelet-TC interactions. In contrast, the metastatic progression of B16F10 was significantly reduced in the absence of GPVI (Fig.5B-C). While B16F10 seeding appeared unaffected (1 dpi), metastatic outgrowth was progressively impaired over time, starting from 7dpi (Fig.5B-C), as we observed in the longTCP model (Fig.4G-H). Macroscopic analysis of lungs revealed again that the number of foci was similar in both conditions, while their size (and proliferation index) was significantly reduced in absence of GPVI (Fig.5D,E). In addition, the number of intrametastatic platelets found was significantly reduced in GPVI-/- mice (Fig.5F). Interestingly their activation ability, measured by the number of aggregates found was not affected. Altogether, these results suggest that the platelet receptor GPVI shapes metastatic outgrowth of TCs that do not rely on early intravascular platelet-TC interactions, without impacting their lung seeding abilities. It further emphasizes the need to consider platelet binding efficiencies and molecular targets before designing platelet-targeted anti-metastatic strategies. Nevertheless, they allow us to validate GPVI as an ideal target, as its minimal effect on bleeding and its contribution to late steps of tumor metastasis make it an appealing candidate for testing its anti-metastatic potential in a human-relevant context.

**Fig. 5.**
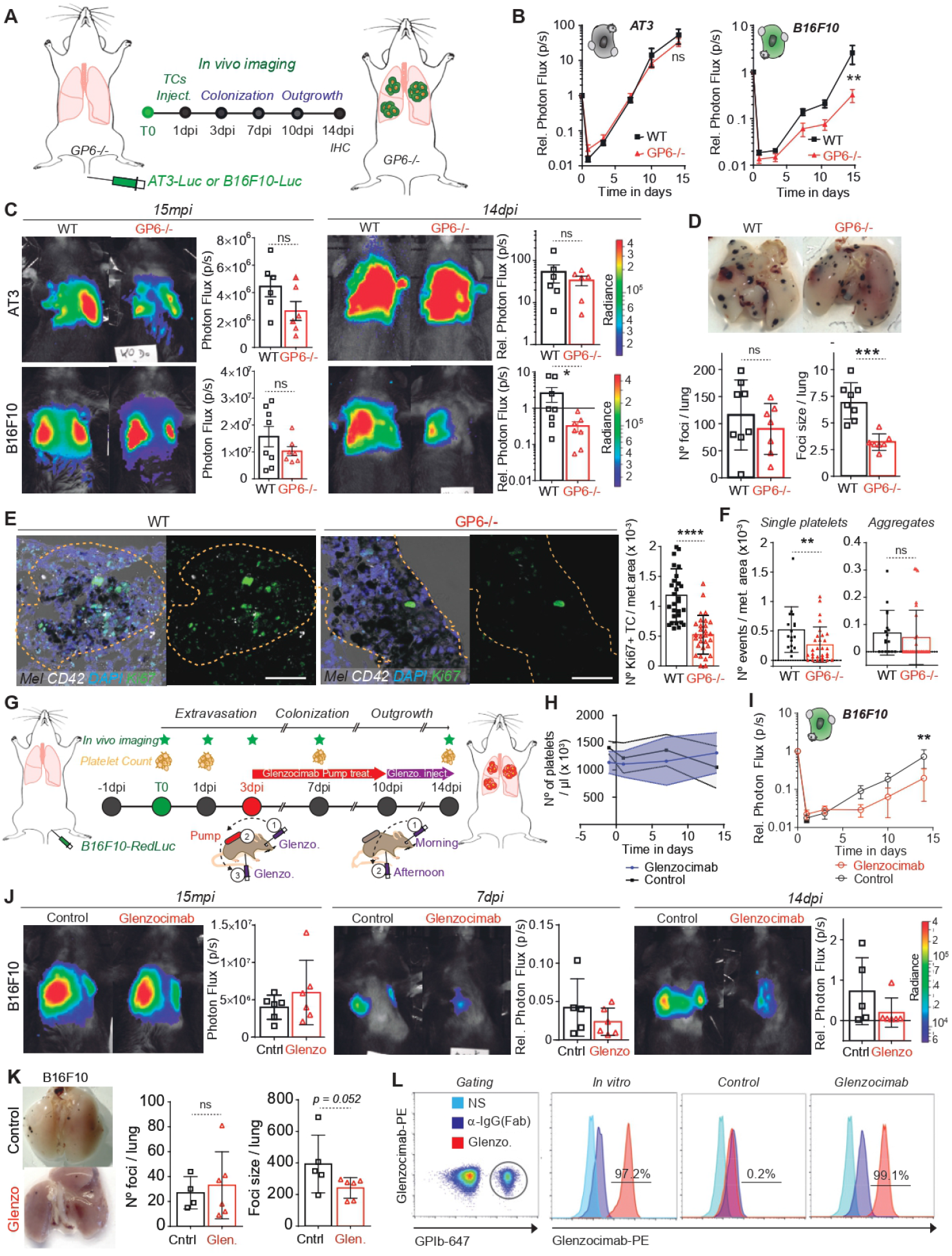
Therapeutic targeting of the human GPVI receptors impairs metastatic outgrowth of established foci. (A) Infographics showing the experimental setting to study the role of GPVI platelet receptor in GPVI-KO mice along 14 days. AT3-Luc or B16F10-Luc TCs are injected intravenously in WT or GPVI-KO mice. Lung seeding and metastatic outgrowth is followed by *in vivo* bioluminescence. (B) Bioluminescence kinetics of AT3-RedFluc-L2 and B16F10-Luc TCs lung outgrowth in WT and GPVI-KO mice. Data are presented as mean +/- SEM. (C) Representative images and time point comparisons (15mpi, left; 14dpi, right) of total lung bioluminescence in normal versus GPVI-depleted animals in AT3 (up) and B16F10 (down) TCs models. Data are presented as the mean +/- SEM, Mann-Whitney analysis. (D) Macroscopic analysis of mice lungs injected with B16F10 cells at 14dpi. Pictures of normal (left) and GPVI-deficient (right) lungs are show. Graphs show the results of B16F10 foci number (bottom left) and size (bottom right) analysis performed on ImageJ. Data are represented as mean +/- SD of 8 mice per group from 1 experiment. (E) Representative zoomed images of the immunohistochemical analysis of lung tissue sections containing B16F10 metastatic foci. Scale =50µm. Proliferative cells within metastatic foci are labelled using the Ki67 marker (green) and platelets with an α-CD42b antibody (white). Graph data (right) are presented as the mean +/- SEM. (F) Graph shows the number of intrametastatic platelets per metastatic area in control and GPVI-KO animals. Data are presented as mean +/- SD of 10 or more images per mice, from 2 to 3 mice and from 1 experiment. (G) Infographics showing the experimental setting to study the impact of GPVI blockade in the outgrowth of B16F10 lung metastasis. B16F10-Luc TCs are injected intravenously in control or α-hGPVI Fab-treated mice (Glenzocimab). Due to the short half-life of the Glenzocimab in mice, the administration is performed by a constant delivery via subcutaneous osmotic pumps. The pumps are implanted 3 days post TCs injection to ensure the establishment of micro-metastatic foci. In this setting TCs outgrowth is followed by *in vivo* bioluminescence and thenanalysis of lung metastatic foci is performed by macroscopic inspection at 14dpi. (H) Graph showing platelet counts on control and Glenzocimab-treated animals. (I) Bioluminescence kinetics of B16F10-Luc TCs lung outgrowth in normal and short-term TCP mice. Data are presented as mean +/- SEM, 2 independent experiments of n=3. (J) Representative images and time point (15mpi, 7dpi and 14dpi) comparisons of total lung bioluminescence in control versus Glenzocimab-treated animals. Data are represented as mean +/- SEM (K) Macroscopic analysis of mice lungs at 14dpi. Pictures of control (top) and α-hGPVI-treated (bottom) lungs with B16F10 metastatic foci are shown. Graphs show the results of B16F10 foci number (middle) and size (right). Data are represented as mean +/- SD. (L) Flow cytometry analysis of mouse platelets labelled *in vitro* and *in vivo*. First histogram refers to the electronic gating strategy on platelets labelled with α-GPIb-647. Second histogram refers to platelets stained with Glenzocimab plus an anti-human (Fab) –PE-labelled antibody *in vitro*. Two last histograms refer to mouse platelets from control or Glenzocimab-treated animals stained with the anti-human (Fab) – PE-labelled antibody.

### Targeting the human GPVI receptor with a humanized antibody impairs metastasis

While this study and others (43) confirm that GPVI is a promising target for impairing metastasis, its relevance to human pathology is missing. Thus, better preclinical models and humanized therapeutic tools, with ideally no thrombocytopenic effects, are needed. To do so, we exploited our previously established genetically modified mouse strain expressing human GPVI (hGPVI) (45). While these mice are viable with no hematological defects, they express a knock-in copy version of hGPVI that makes it the best candidate to test anti-GPVI compounds. To counteract GPVI function, we decided to exploit Glenzocimab (previously ACT017, Acticor Biotech (46). Glenzocimab is a humanized antibody fragment (Fab) that has recently completed a phase II trial in stroke patients and is currently examined in a phase II/III trial (ACTISAVE, ClinicalTrials.gov Identifier: NCT05070260). This compound binds to hGPVI to block ligand binding, which occurs without causing thrombocytopenia or GPVI membrane depletion, and is not associated with bleeding events (47).

Building on the exciting observation that lateTCP could reduce the growth of established metastases (Fig.4), we subjected hGPVI mice to an experimental metastasis protocol targeting the growth of established metastases (mimicking lateTCP, Fig.5G). The half-life of Glenzocimab in mice is rather short (data not shown) when compared to humans (47). We thus administrated Glenzocimab progressively using osmotic pumps that were implanted subcutaneously (Fig.5G). We confirmed that Glenzocimab did not affect platelet counts (Fig.5H) and was found to be successfully bound to mouse platelets (Fig.5L) four days after pump implantation. Longitudinal imaging of B16F10 lung metastasis revealed that the metastatic burden of hGPVI mice treated with Glenzocimab was significantly reduced starting from 4 days after the implantation of the pump (Fig.5I,J). Metastasis growth was constantly reduced, reaching a 4-fold decrease at 14 dpi (Fig.5I). Macroscopic inspection of the lungs revealed that Glenzocimab had, as expected, no effect on the number of metastatic foci, but reduced their size (Fig.5K). Altogether, these results identify, for the first time, an anti-GPVI compound that can target human GPVI and impair metastatic progression with no side effects (thrombocytopenia, bleeding). Considering the very positive safety profile of Glenzocimab and its efficiency in repressing growing metastasis, we provide here a new therapeutic agent with previously unmet clinical potential.

## DISCUSSION

This manuscript reports the first longitudinal analysis on the pro-metastatic role of platelets in lung metastasis by subjecting two TC models (breast carcinoma and melanoma) to experimental metastasis assays in the zebrafish embryo and mice. These were tested in six mouse models of time-controlled thrombocytopenia and by the genetic and pharmacological ablation of the platelet receptor GPVI. By these means, we have discovered that different TCs interact with platelets differently, and confirmed that such ability shapes intravascular arrest and metastatic fitness of TCs. Importantly, we provide the first study demonstrating that platelets contribute to late steps of metastasis and document the therapeutic efficacy of targeting GPVI in animals carrying growing metastases of TCs with low platelet binding potential. Altogether, our data identify GPVI as the first molecular target whose inhibition can impair metastasis without inducing collateral hemostatic perturbations.

The established mechanisms proposed for the pro-metastatic role of platelets including platelet binding, activation, and aggregation by TCs, have been traditionally insufficient in providing momentum for the development and establishment of anti-platelet drugs in routine oncologic care. This may lie in the exclusive focus in a too narrow and early window within the metastatic process, thus leaving few options for therapeutic intervention. Current data assume that all cancers interact equivalently with platelets, which lead to controversial results in different cancer models (25,26) and unclear outcomes when using anti-platelet drugs in oncologic patients (27,48). For example, thrombocytosis is a poor prognosis factor, reflecting an aggressive tumor phenotype in cervical, colon, and non-small cell lung cancer patients (49,50). On the contrary, in pancreatic cancer, thrombocytopenia was associated with a worse prognosis, thus indicating the varied influence of platelets depending on the type of cancer (51,52). While CTCs are widely considered to be covered by platelets when entering the blood (20), whether this applies to all cancer types remains to be demonstrated. Here, we show that TCs do not all bind platelets equally (Fig.1) and importantly, that TCIPA seems to occur only at high concentrations of TCs *in vitro* (28,53,54), which is an unlikely scenario *in vivo*. All these issues have left anti-platelet drugs out of the standard cancer treatments and/or prophylaxis. Aware of this, and the lack of longitudinal studies of the whole lung metastatic cascade *in vivo*, we developed a novel experimental approach merging the single cell analysis of *in vivo* arrested TCs combined with *in vivo* bioluminescence longitudinal analysis of micro- and macro-metastases in mouse models of thrombocytopenia. This approach allowed us to identify a novel prometastatic role of platelets in the late stages of the disease mediated by the GPVI platelet receptor and allowed us to probe the feasibility of targeting it with a therapeutic antibody.

First, we identified differential platelet binding profiles in different TC types. We are aware that our binding results may not mimic physiological conditions, but this simple approach proved to be a good starting point to enquire about the specific interaction of platelets and different TCs in more complex models integrating blood flow forces. Indeed, static binding profiles correlated with the *in vivo* prometastatic effects observed in both ZF and mouse, although their absolute binding efficiencies differed. It has been shown that hydrodynamic shear affects the receptor specificity of activation-dependent platelet binding to TCs, as evidenced by the transition from a P-selectin-independent/Arg-Gly-Asp (RGD)-dependent process at 100 s-1 to a P-selectin/αIIbβ3- dependent interaction at 800 s-1 (55). This may imply that static assays such as SEM or aggregations are overestimating platelet binding, as in such a scenario resting platelets could bind independently of P-selectin and αIIbβ3 integrin. On the contrary, in an *in vivo* situation platelets must be activated to bind TCs. Vascular regions with low flow profiles may thus increase platelet binding through this P-selectin/αIIbβ3 independent mechanism. Interestingly, platelets are most prometastatic in the cell model able to bind more platelets in static conditions (4T1 cells), meaning that low flow binding abilities are capital for tuning TC’s metastatic fitness. Using our zebrafish model, we also observed that platelets favor the stable adhesion of 4T1 cells exclusively, indicating that an early and active platelet binding is capital for the arrest of TCs in blood vessels that aren’t physically constraining to TCs. The putative molecular players involved in this differential binding remain to be investigated.

To further understand the mechanisms driving the prometastatic effect of platelets at early stages, we performed an exhaustive analysis by quantitative confocal imaging of single cells arrested in mouse lungs. This allowed us to characterize the dynamics of the arresting site and probe TCs viability and platelets recruitment. We first confirmed that binding profiles (not efficiencies) found *in vitro* in static conditions are conserved *in vivo*. However, the platelet binding profile changed only 2 hours post-injection, with B16F10 cells now attracting increasing numbers of platelets. In this sense, we found that platelets favour the survival of intravascularly arrested TCs by direct binding to the surface of viable cells or by passive recruitment around non-viable cells. This passive recruitment may occur by vessel occlusion or release of platelet-activating cytoplasmic content such as tissue factor (TF) (22,56). In the first case, active recruitment leads to a highly significant long-term metastatic fitness enhancement (Fig.3), while in the latter case, passive recruitment leads to a discrete but significant metastatic fitness enhancement, only appreciated at late time points (14dpi). Previous work has suggested a key role for thrombin and clot formation as the starting point of brain metastasis (22), however, not all metastatic cell lines express TF, the trigger of thrombin generation. Considering that metastasis is a very inefficient process and that intravascular cell death is a well-known cause for TF release and thrombin production, the co-option of viable with non-viable cells may help the viable cell to successfully metastasize, as previously observed (57). Anyhow, this difference between active and passive platelet recruitment mechanisms may be capital for designing anti-platelet drugs for treating different cancer types.

Previous reports have found evidence of platelets inside primary tumors (14,34,35) with no evidence that this could apply to metastasis. We now document that metastatic sites are populated by platelets suggesting that they might be able to extravasate with or independently of TCs, as reported previously (58). Platelets have been shown to migrate easily within tissues (59) suggesting that they could actively populate growing metastases. Whether platelets require extravasation to seed metastases remains to be demonstrated, as platelets might just interact and imprint TCs in circulation. For example, while TCs interact with platelets, fibrin, or fibrinogen from 0 to 6h post-injection, platelets are not found at intravascular arresting sites from 6 to 24 hpi (52), meaning that early intravascular interactions are frugal. In this sense, the GPVI receptor may tune metastasis by passive and intermittent interaction with TCs at the arrest sites (42), similarly as shown by our genetic and pharmacological loss-of-function models at late steps of the cascade.

Whether platelets actively migrate and exchange material within metastatic foci remains to be demonstrated and opens exciting avenues of research. The observation that only the number of platelets but not their aggregation capacity is affected in GPVI-KO animals (Fig.5F) may indicate a defect in migration. It further confirms the safe thrombostatic profile of GPVI-targeted platelets *in vivo* at the cellular level. Platelets are known to release significant amounts of bioactive material within releasates (60) that shape, for example, their invasion abilities. Platelets also secrete microparticles and extracellular vesicles, which are angiogenic (61), immunosuppressive (13) and enhance TCs survival and outgrowth capacity (62), as confirmed in our models. Indeed, we have demonstrated an increased proliferative index in the presence of platelets in our short and lateTCP models. We further observed a platelet-dependent recruitment of immune cells (CD45+) within growing metastatic foci (lateTCP). Besides the early physical protection from immune cells during circulation (shielding), platelets also support the generation of immunosuppressive tumor microenvironments and modulate immune responses broadly. They secrete and express TGFβ, IL-1, PDGF, CCR5, CD40, CD154, and toll-like receptors and can uptake, process, and present antigens to CD8+ T-cells, suggesting that they act as bona fide immune cells (63,64). Additionally, the ratio between lymphocytes and platelets (PLR) has been recently identified as a predictive marker in cancer progression in clinical settings (65-67). Based on these evidences, it is tempting to speculate that platelets may influence metastatic outgrowth by regulating immune surveillance at the tumor microenvironment. For example, they could recruit immunosuppressive populations such as tumor-associated macrophages (TAMs), myeloid-derived suppressor cells (MDSCs, N2 neutrophils and/or tolerogenic dendritic cells (tDCs) (68). However, a deeper analysis characterizing the nature and functional phenotype of immune cell infiltrates in TCP models would help to clarify the platelet-mediated immune responses within the tumor microenvironment, and provide clues as to what triggers metastatic impairment when targeting GPVI.

While reducing platelet count is not a viable therapeutic approach, due to the risk of bleeding, targeting platelet receptors could circumvent this issue. When depleting or targeting GPVI, we could indeed successfully impair the proliferation rate within B16F10 metastatic foci. This points to GPVI as a promising target to develop new anti-metastatic drugs able to target the disease once the metastatic spread is already initiated, which account most of the cases in clinical practice. Such achievement is of utmost importance. We provide here evidence that a pharmacologic blockade of GPVI with Glenzocimab (Acticor Biotech), an anti-GPVI F(ab) fragment evaluated in phase I clinical trials to prevent ischaemic stroke, efficiently reduces experimental metastasis. This drug is particularly promising as it has an anti-thrombotic effect without inducing any bleeding in healthy volunteers and in patients (69). While there are several anti-platelet drugs available in the market, most of them increase the risk of hemorrhage in clinical settings, or have limited efficacy against metastasis, with the exception of aspirin (70). Altogether, our work demonstrates that targeting platelets in animals carrying growing metastases can impair their growth and it further identifies GPVI as the first molecular target whose inhibition can hamper metastasis without inducing collateral hemostatic perturbations.

## ACKNOWLEDGMENTS

We thank all members of JGG’s team for their constant discussions on this particular topic. JGG is the coordinator of the NANOTUMOR Consortium, a program from ITMO Cancer of AVIESAN (Alliance Nationale pour les Sciences de la Vie et de la Santé, National Alliance for Life Sciences & Health) within the framework of the Cancer Plan (France). Work and people in the lab of JGG are mostly supported by the INCa (Institut National Du Cancer, French National Cancer Institute), charities (La Ligue contre le Cancer, ARC (Association pour la Recherche contre le Cancer), FRM (Fondation pour la Recherche Médicale)), the National Plan Cancer initiative, the Region Grand Est, INSERM and the University of Strasbourg. This work has been directly funded by the INCa grant (PLBIO 2014-01) and by the support of the Ligue contre le Cancer (labelisation) and the SATT Conectus (Strasbourg). MGL has been funded by the University of Strasbourg (IdeX, investissements d’Avenir), the INCa and the SATT Conectus (Strasbourg). MP is supported by INCa, FC by NANOTUMOR and VM by a Ph.D. fellowship from the French Ministry of Science (MESRI). GF was supported by La Ligue contre le Cancer. We are grateful to Acticor for providing the Glenzocimab and to François Bertucci for providing human samples. We are also thankful for recent donators (Rohan Athlétisme Saverne) to support our work.

## ETHICS STATEMENT

Human studies were performed according to Helsinki declaration. Control human samples were obtained from volunteer blood donors who gave written informed consent recruited by the blood transfusion center where the research was performed (Etablissement Français du Sang, Grand-Est). All procedures were registered and approved by the French Ministry of Higher Education and Research. The donors gave their approval in the CODHECO number AC-2008 - 562 consent form, for the samples to be used for research purposes. Immunocompetent and genetically modified mice used in this study were housed under pathogen-free conditions and all procedures were performed in accordance with the European Union Guideline 2010/63/EU. The study was approved by the Regional Ethical Committee for Animal Experimentation of Strasbourg, CREMEAS (CEEA 35) and registered under the APAFIS authorization 14741-2018041816337540.

## MATERIAL and METHODS

### Cell lines

The mouse breast cancer cell lines 4T1, E0771, AT3, and D2A1, and the mouse melanoma cell line B16F0, were cultured under standard conditions (37°C, 5% CO2) using RPMI-1640 or DMEM, respectively, supplemented with 10% FBS and 1% penicillin-streptomycin solution. The human breast cancer MDA-MB-231 and melanoma cell lines A375 and MCF7, we cultured under the same conditions (DMEM). Their viabilities *in vitro* were assayed before aggregation and SEM assays and *in vivo* experimental metastasis experiments by an ADAM-MC Automated cell counter (NanoEntek).

### Cell line engineering

For *in vivo* imaging on zebrafish embryos, 4T1 and B16F10 cells were engineered to express a Lifeactin-tdTomato fusion protein. Briefly, the Life-tdTomato DNA fragment derived from the Addgene plasmid tdTomato-Lifeact-7 (number 54528) was inserted in the pLSFFV-Ires-Puromycin or pLenti-CMV-mPGK-Puromycin lentiviral vectors to generate the pLSFFV-LifeActin-tdTomato-Ires-Puro and the pCMV-LifeActin-tdTomato-mPGK-Puro lentivirus lentiviral vectors used to transduce 4T1 and B16F10, respectively. Lentivirus particles production and transduction procedures have been already described in (32). For confocal imaging of single arrested TCs in mice lungs, luciferase-expressing 4T1 or B16F10 cells were labeled *in vitro* with CellTrace Yellow (Thermo Fischer) according to manufactured instructions.

For *in vivo* imaging in mice, regular firefly luciferase or red-shifted firefly luciferase (Bioware® Brite B16F10-Red-FLuc, Perkin Elmer) was used. B16F10 cells expressing red-shifted luciferase were purchased from Perkin Elmer. 4T1 luciferase-expressing cells were a kind gift from Corinne Laplace-Builhé (Institut Gustave Roussy Paris). To generate a luciferase-expressing breast cancer AT3 cell line (44), we engineered it to express the red-shifted luciferase protein using the RediFect Red-Fluc Lentiviral Particles from Perkin-Elmer. To avoid immune rejection in immunocompetent C5BL6/J recipients, an *in vivo*-derived AT3 cell line was established from lung metastases derived from a tail vein-injected mouse. Briefly, mouse lungs containing AT3-RedFLuc metastasis were collected on Day 18 post-injection and chopped into 1 mm3 pieces with a scalpel. The pieces were resuspended in 4000µl of dissociation buffer (Miltenyi Biotec Neural Tissue Dissociation Kit (P) 130-092-628) in the presence of 100µl of collagenase solution (Sigma-Aldrich C9891) at 100mg/ml together with 400µl of DNaseI at 1mg/ml under gentle mixing at 37°C during 10 min. 50µl of enzyme P was added and further incubated for 5 min. 60µl of enzyme A was added with an additional incubation period of 5 min. The suspension was then ground on a Corning cell strainer of 40µm (reference 431750) over a 50ml falcon tube using a 2.5ml syringe rubber plunger. Four ml of full cell medium were added to the filtrated solution and cells were then collected by centrifugation, 5 min, 400g at room temperature. The pellet was resuspended in DMEM medium containing 10% FCS, 1% PS, 1% Gentamicin, 5µg/ml of Fungizone, and puromycin at 1µg/ml in a T75 flask. After a few days of puromycin selection, the cell line was considered established and called AT3-RedFluc-LM1. To ensure the absence of immune rejection on recipients a second round of *in vivo* derivation was performed, thus generating the AT3-RedFluc-LM2 cell line which was used in our experiments.

### Platelets isolation

For obtaining human citrated platelet-rich plasma (cPRP) blood was collected in citrate buffer (3.8%). For obtaining washed human platelets suspensions, the blood was collected in the presence of 1/7 volume of citric acid-citrate-dextrose (ACD) pH 6.5, from healthy volunteers with informed consent, who had not taken any drugs known to affect platelet functions for at least 10 days before the study. Citrated platelet-rich plasma was prepared by centrifugation at 250g for 16 min at room temperature and collection of the supernatant. Washed platelets were prepared as described previously (71). Briefly, blood was centrifuged at 250g for 16 minutes at 37C. Supernatant (PRP) was collected and centrifuged at 2200g (time of centrifugation depends on volume). Platelet pellet was resuspended in pre-warmed (37C) Tyrode-Albumin buffer (TA) (0.35% HAS, glucose 0.1%, NaCl 137 mM, KCl 2.7 mM, NaHCO3 12 mM, NaH2PO4 0.36 mM, HEPES 5mM, CaCl2 2 mM, MgCl2 1 mM, pH 7.35, 295 mOsm) containing 10U/mL of Heparin and 0.5 µM of PGI2. Platelet suspension was incubated for 10 minutes at 37C. PGI2 was added again to the tubes, centrifuged at 1900g for 8 minutes and the supernatant was discharged. Pellet was re-suspended in TA buffer containing 0.5 µM of PGI2 and centrifuged again after 10 minutes incubation at 37C. Pellet was re-suspended in TA buffer with 0.02U/ml of apyrase at a platelet concentration of 300 × 1e3 platelets/µl. The quality of the washed platelet suspension was routinely tested by adding ADP 5µM as agonist in a light aggregometry assay in the presence of fibrinogen, or in its absence, in the case of cPRP suspensions. For obtaining mouse washed platelet suspensions, the blood was collected by an aortic puncture after intraperitoneal injection of anesthesia (ketamine 100mg/kg, xylazine 20mg/kg). Anesthetized animals were placed on heating pads at 38 C for dissection. By the use of a sterile tissue pad, abdominal organs were side displaced to show the abdominal portion of the aortic artery. A preloaded syringe with ACD was used to collect the blood by punction at the level of the separation of the aortic artery into the two iliac arteries. Around 700-800 µl of blood were collected per mouse. Mice were sacrificed while anesthetized by cervical dislocation. For obtaining mouse citrated platelet-rich plasma (cPRP) blood was collected by aortic puncture in citrate buffer 0.315% following centrifugation of the whole blood in microtubes during 1 min at 1900g at room temperature.

### Zebrafish handling and intravascular injection of tumor cells

*Tg(fli1a:eGFP)* Zebrafish (Danio rerio) embryos from a Tubingen background used in the experiments were kindly provided by the group of F. Peri from EMBL (Heidelberg, Germany) and further grown and bred in our in-house Zebrafish facility. Embryos were maintained in Danieau 0.3X medium (17,4 mM NaCl, 0,2 mM KCl, 0,1 mM MgSO4, 0,2 mM Ca(NO3)2) buffered with HEPES 0,15 mM (pH = 7.6), supplemented with 200 µM of 1-Phenyl-2-thiourea (Sigma-Aldrich) to inhibit the melanogenesis, as previously described (72). Forty-eight-hour post-fertilization (hpf) *Tg(Fli1a:eGFP)* embryos were mounted in a 0.8% low melting point agarose pad containing 650µM of tricaine (ethyl-3-aminobenzoatemethanesulfonate) to immobilize the embryos. TCs LifeAct-TdTomato cells were injected with a Nanoject microinjector 2 (Drummond) and micro-forged glass capillaries (25 to 30 µm inner diameter) filled with mineral oil (Sigma). For the intravascular injection of TCs in the zebrafish embryo vasculature, previously washed (EDTA 0.48 mM, Versene, Gibco) and gently detached (Typysin solution 0.125% diluted in Versene 1x, Gibco) 4T1 or B16F0 TCs were washed and resuspended in serum-free RPMI-1640 or DMEM media at a concentration of 100 × 1e6 TCs/mL and kept on ice until injection. Prior injection 50µl of TCs suspension were added to 50µl of human cPRP at a platelet concentration ranging from 350 to 450.000 platelets/µL (1/4 ratio aprox.) and incubated at 37C during 2-3 minutes. Human citrated platelet poor plasma (cPPP) was used as a control (no platelets condition). Finally, 23nL of the TC-platelets mix was then injected in the duct of Cuvier of the embryos under the M205 FA stereomicroscope (Leica), as previously described (73).

### Mice

Male WT C5BL6/J immunocompetent mice from 8 to 16 weeks were used for the B16F10 melanoma cell model, while female WT immunocompetent Balb/c mice were used for the 4T1 breast cancer model. All were purchased from Charles River Labs. C5BL6/J immunocompetent mice deficient for the platelet receptor GPVI (GPVI-/-), and bearing the human form (hGPVI) were generated as previously described (45); males and females were used from both models. Animals were housed in pathogen-free conditions. Mice received standard chow and sterile tap water ad libitum. Appropriate enrichment (sterile pulp paper and coarsely litter) was provided. Physical condition of mice was monitored daily. A protocol approved for early euthanasia of terminally sick animals was in place. All animal procedures were performed in accordance with institutional guidelines and approval, under the APAFIS authorization 14741-2018041816337540

### Platelet and GPVI receptor depletion and pharmacological blockage

Severe thrombocytopenia was induced by intravenous (i.v.) injection of 50µg (2mg/kg) of in-house-produced αGPIb antibody (74) or IgG isotype control per mouse (IgG, 1mg/mL; RAM.6, 2 mg/mL). Injections were delivered at precise time points to differently affect the metastatic cascade, meaning 24 hours prior TCs injection, 3dpi, 7dpi and/or 10dpi. To specifically deplete GPVI receptor in mouse platelets, 50µg of JAQ1 mAb (1mg/mL, Emfret) was injected subcutaneously per mouse. Control animals were injected with 50µg rat IgG isotype control (Emfret). For the delivery of anti-human GPVI antibody Glenzocimab, due to its short half-life in the mouse circulation, an Azlet osmotic pump (model 2001) was implanted subcutaneously (s.c.) on the dorsal side, to deliver the antibody for 7 days, at a rate of 1µl/hours at a concentration of 20mg/ml.

### Platelet Counts

Murine whole blood was collected into EDTA (6 mM) after severing the mouse tail. The platelet count and size were determined in a Scil Vet abc automatic cell counter (Scil Animal Care Company, Holtzheim, France) set to murine parameters. The platelet count on human platelet suspensions (cPRP and/or washed platelets) was assessed in an automatic platelet counter (XN-1000, Sysmex).

### Experimental pulmonary metastasis assay and bioluminescent imagings

Subconfluent 4T1 and B16F10 cells expressing regular luciferase (Photinus pyralis) or its red-shifted version, respectively, were washed with EDTA 0.48 mM (Versene, Gibco), gently detached using a 0.25% trypsin-0.02% EDTA solution (Gibco), washed in media containing 10% fetal calf serum, resuspended at a concentration of 1.5 × 1e6 TCs/mL in serum-free media, filtered through a 40 µm mesh (filcon) and kept on ice until injection. Viability was determined by trypan blue exclusion and was always more than 85% prior injection. About 100 µl of TCs suspension were injected per mouse through the lateral tail vein by using a 25-gauge needle. *In vivo* imaging of mice lungs was immediately performed after TCs inoculation to establish initial lung seeding (15 mpi). Subsequent imaging points at 0.5, 1, 2, 4 and 24h, and 1, 3, 7, 10 and 14 days after TCs injection were monitored. To do so, 5 min after intraperitoneal injection of D-luciferin solution (150mg/kg) to the isofluorane-anesthetized (Isoflo, Zeotis) mouse, the ventral bioluminescence image of mice lungs was acquired with an IVIS Lumina III (Perkin Elmer) imaging system and then analysed using the Living Image software (Perkin Elmer). The rate of total light emission of the lung metastatic area was calculated and expressed as numbers of photons emitted per second (p/s).

### Light transmission aggregometry (LTA)

TC-induced platelet aggregation (TCIPA) was analyzed by LTA. Human citrated platelet rich plasma (cPRP) and washed platelets were isolated from citrated or ACD blood of human donors, respectively, according to the approval of the local ethics committee. Mouse citrated PRP was obtained as previously described. The aggregation procedure was performed equivalently for human or mouse samples. Briefly, platelet aggregation was measured at 37°C by a standard turbidimetric method in an APACT 4004 aggregometer (ELITech Group, Puteaux, France). Serum free media or TA buffer were used as control together with PEG-treated 10µm polystyrene beads (Phosphorex). Aggregations were launched in aggregation vials by adding 200µl of washed platelets at a final platelet concentration of 3e5/µl, or non-diluted citrate platelet rich plasma (cPRP). Next, 50µl of TCs (or control beads) resuspended in TA buffer or serum free culture media were added (final concentration up to 3e6/ml). Platelets suspensions were stirred at 1,100rpm. Control samples were activated by the addition of 5µM ADP in the presence or absence of human fibrinogen (320 µg/mL, in-house generated (71)) or native equine tendon collagen Type 1 (2.5 µg/mL, Collagen Reagens HORM® Suspension, Takeda) in a final volume of 300µL. Prior aggregation, TCs were washed and gently detached by non-enzymatic means (EDTA 0.48 mM, Versene 1x, Gibco), washed twice in RPMI-1640 or DMEM medium without serum + 0.1% BSA + 2mM CaCl2 + 1mM MgCl2, or in TA buffer. Platelet aggregation was evaluated by comparing the amplitudes of the aggregation curves generated with APACT LPC software. The initial drop in light scattering due to sudden TC addition to running aggregation vials was subtracted in Microsoft Excel to obtain adjusted and comparable aggregation curves.

### Scanning Electron Microscopy

Gently detatched TCs (EDTA 0.48 mM, Versene, Gibco) were washed in serum free media (cPRP) or TA buffer (washed platelets) prior incubation with platelet suspensions at the previously indicated concentrations. PEG (Polyethylene glycol) blocked beads were used as a control. TCs and platelet mixes resulting after LTA assay (300 µl) were immediately fixed in 300 µl of 2x fixative solution (4% paraformaldehyde, 5% glutaraldehyde in standard 0.1M Sodium Cacodylate buffer, NaCac). TCs and platelet aggregates were sedimented O/N at 4ºC. Supernatants were removed and pellets gently resuspended in NaCac Buffer. TCs and platelets suspensions were placed on p24 well plates containing glass coverslips previously treated with poly-L-lysine solution (Sigma). After TC-platelet suspension sedimentation (30 minutes at room temperature) on the coverslip, supernatant was removed and coverslips fully dried in a 60C oven during 1h. Wash coverslips with 0.2 um-filtered NaCac buffer and dH2O and start sample dehydration in graded Ethanol series (1 × 70%, 1 × 80%, 1 × 95% during 5 min each and 2 × 100% during 30 minutes each). For final and irreversible sample dehydration a graded serie (25%, 50%, 75% and 100%, 5 min each) of 1,1,1,3,3,3-hexamethyldizilazane (HDMS, Merck, Millipore) was used. Cells were mounted onto 12 mm EM Aluminum Mounts for AMRAY (EMS, Hatfield) by using Leit-C conductive carbon cement (CCC, Plano GmbH, Germany). Platinum metallization of the sample was performed under vacuum by using a Cressington Sputter Coater 208HR coupled to a Pfeiffer Vacuum (Germany). Image acquisition was performed at high-resolution (10-15kV) and 6000-10000x magnification on a Phenom-World SEM desktop microscope (Phenom-World B.V, The Netherlands).

### CLEM sample preparation and image acquisition

Correlative Light and Electron Microscopy (CLEM) was performed as previously described (72). To characterize ultrastructural features of CTC-bound platelets interacting with the ZF endothelium, chosen ZF embryos were chemically fixed with 2.5% glutaraldehyde and 2% paraformaldehyde in 0.1M NaCac buffer (pH 7.4). The sample was kept in fixative for 48hr at 4°C and washed 3 times with 0.1M NaCac buffer, pH7.4, followed by a second fixation with 0,05% malachite green, 2.5% glutaraldehyde in 0.1M NaCac buffer, pH7.4 during 25 min in an ice bath. The sample then was post-fixed for 45 min in 1% OsO4 in 0.1M NaCac buffer, pH7.4 in a fume hood in an ice bath and washed 2 times with in 0.1M NaCac, (the buffer was also kept in an ice bath to avoid thermal changes). Then the specimen was incubated in 1% aqueous tannic acid solution for 25 min in an ice bath and finally washed 5 times with distilled water (DW). The zebrafish sample was dehydrated with serial ethanol solutions (25%, 50%, 70% 95% and 100%) and dry acetone at RT for 10 min. Subsequently the sample was incubated in a serial resin-acetone solution (1:3; 1:1; 3:1). Further, the sample was incubated in 100% Epon resin 3 times 1h at RT. The sample was allowed to polymerize in an oven at 60°C for 48h. The resin block was trimmed by ultramicrotomy. After targeting region of interest (ROI), 90 nm thin sections were collected and placed in slot Formvar cupper grids. The TEM data set was acquired with a Hitachi 7500 TEM, with 80 kV beam voltage, and the 8-bit images were obtained with a Hamamatsu camera C4742-51-12NR. The collected images were processed for segmentation using an open-source software ImageJ; AMIRA and IMARIS as follows: a) single images were stitched in a bigger composed image and combined into a 3D-stack; b) The combined 3D-stack was aligned and the contrast adjusted, by ImageJ. C) The segmentation of objects of interest, (arrested tumor cells, platelets) and, 3D Volume reconstruction, by AMIRA and IMARIS.

### Tissue histology and immunohistochemistry

Human melanoma samples from lung metastases were obtained, processed and stained as previously described (75). For murine samples, mice were sacrificed under isofluorane anesthesia and lidocaine analgesia (5mg/kg) at 15mpi, 2hpi and 15dpi. Lungs were resected, washed and fixed O/N in 4% paraformaldehyde/phosphate buffered saline (PBS) solution (EMS). Left lung lobes were then treated for paraffin inclusion, while the right lung lobes were treated for Optimal Cutting Temperature (OCT) inclusion (Tissue freezing media, Leica). Briefly, O/N fixed lungs were subjected to dehydration on serial grades of ethanol (1 × 50%, 1 × 80%, 1 × 95% and 2 × 100% during 1 hour each) and a final xylene/toluene bath (2 × 45 min each). Dehydrated tissue was then paraffin embedded in an automated Leica inclusion machine during 2 hours. Cryosections of 20µm (for single cell analyses and segmentation) or 10µm (for regular IF analyses) paraffin sections were mounted on poly-lysine-coated slides (SuperFrost UltraPlus, Thermo Fisher Scientific). In the case of paraffin-embedded samples, tissue antigens were retrieved by boiling in sodium citrate solution (10 mM, pH 6.0, Sigma). For blocking unspecific antibody binding sites, samples were incubated for 1 h in blocking solution (3% bovine serum albumin, 20 mM MgCl2, 0.3% Tween 20, 5% fetal bovine serum in PBS). Background and nonspecific staining was determined by incubating tissue samples in the absence of primary antibody. Proliferation index within metastatic foci was calculated by the staining with a rabbit anti-mouse Ki67 antibody (clone SP6, Thermo Fisher). For CD45 staining, a mouse biotinilated anti-mouse CD45 antibody (clone 30-F11, Biolegend) was used. Prior to the addition of secondary antibody, tissue endogenous biotin was quenched with Avidin/Biotin blocking solution (Vector Laboratories). Development of the biotinilated antibody signal was done by 1 h incubation with the Avidin/Biotin-HRP complex (Elite Vectastain ABComplex Kit, Vector Laboratories) following 1 h incubation with Streptavidin-647 (BioLegend). For metastases delimitation cytokeratin (mouse anti-mouse cytokeratin pan-mixture, ascites, Sigma) and luciferase (rabbit anti-firefly luciferase, ab21176, Abcam) expression was used in 4T1 and B16F10 metastatic foci, respectively. For the staining of mouse platelets an in-house-generated and directly labeled (488 or 647) anti-GPIb antibody (Rat anti-mouse GPIb (CD42b) RAM.1) was used. For the detection of primary antibody signals, the following Alexa-labeled (Invitrogen) secondary antibodies were used: goat anti-rabbit (H+L) Highly Cross-Adsorbed Alexa-555, goat anti-mouse (H+L) Highly Cross-Adsorbed Alexa-555, goat anti-rat (H+L) Highly Cross-Adsorbed Alexa-488 or 647. In all cases, nuclei were stained with DAPI (Thermo Fisher Scientific) and slides mounted with Fluoromount-G (SouthernBiotech).

### Laser Scanning Confocal Microscopy (LSCP)

For immunohistochemical analyses images were acquired using a Leica TCS SP8 laser scan confocal microscope using a 20x/1.0 W-Plan-Apochromat objective (oil (NA 0.75) and a 63×Plan-Apochromat [oil (NA 1.4)] magnification. For 3D segmentation of cells and platelet aggregates, no less than 10 z-stacks (1AU) of single cells or clusters were acquired in 12-bit color depth and non-saturating settings.

For *in vivo* imaging of TCs circulation and arrest on the *Tg(fli1a:eGFP)* zebrafish embryos a Leica TCS SP8 microscope equipped using an M205 FA stereomicroscope (Leica) and a 20x/1.0 W-Plan-Apochromat objective (oil (NA 0.75)) on an SP8 confocal microscope (Leica) were used.

### Image processing and analysis

Metastatic foci number and size was obtained by image processing on ImageJ software. Briefly, macroscopic 8-bit lung (B16F10-seeded) images (MacOS 15.4, iPhone8) with identical zooming and size (100 × 100 px) were loaded on imageJ. Automatic threshold of the images yield binary masks that were analyzed by the “analyze particles” plugin. Circularity was set between 0.2-1.0 to exclude lung edge shadows. Pixel size bellow 15px was considered as noise. Platelet aggregates and cell volumes arrested intravascularly in mice lungs were obtained by segmentation in AMIRA Visage 6 after confocal acquisition of Z-stacks covering the whole cell-aggregate volume. For immunofluorescence images brightness and contrast were equally adjusted in control and treated samples on ImageJ and/or Adobe Photoshop CS4 software.

### Statistics

Statistical analysis was performed with the GraphPad Prism 9.4. The normal distribution of the data was tested using the Shapiro–Wilk normality test. When comparing two groups, we used a Mann-Whitney analysis. When more than 2 groups were compared, we used a Kruskal-Wallis test followed by the original FDR method of Benjamini and Hochberg post test. When experiments were analyzed as kinetics, we used a two-way ANOVA followed by the original FDR method of Benjamini and Hochberg post test (*p<0.05; **p<0.01, ***p<0.001, ****p<0.0001). In all cases, the α-level was set at 0.05. All the data in graphs was presented as mean +/- Standard Deviation, except for the kinetics that was presented as mean +/- SEM.

**Fig.S1.6.**
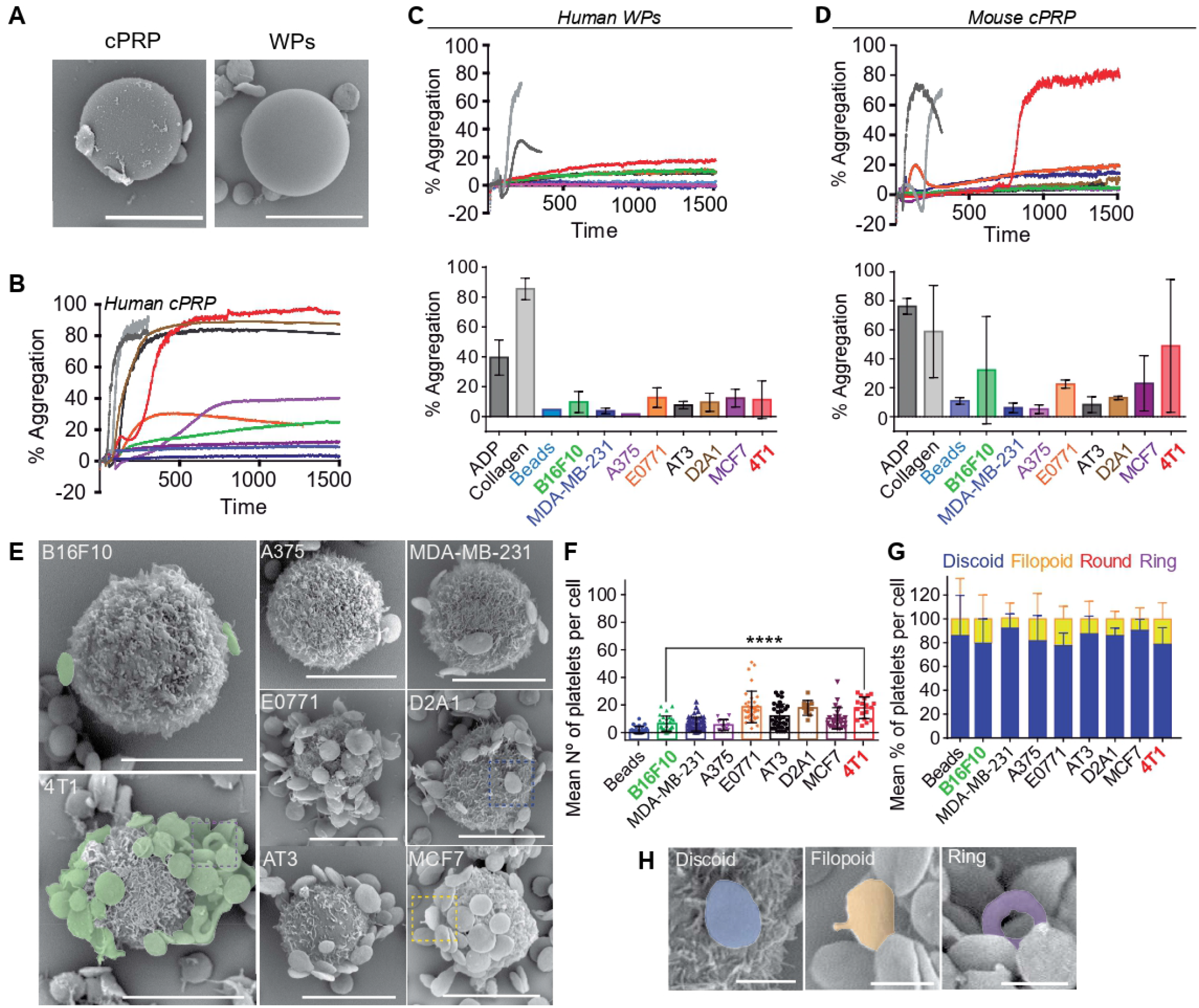
SEM controls and representative aggregation curves. (A) SEM images of PEG-treated 10µm polystyrene beads interacting with mouse platelets contained within mouse citrated platelet rich plasma (cPRP) or human washed platelets (WPs). (B) Representative platelet aggregation curves obtained with human cPRP for each TC line. ADP and collagen were used as positive controls, while beads were used as negative control. (C, D) Representative platelet aggregation curves (up) and platelet aggregation efficiency (%) (down) of the different TCs tested using human WPs (C) or mouse cPRP. No significant differences between 4T1 and B16F10 aggregation efficiency was observed due to the high variability of the assay. Representative data from 2 to 4 independent aggregation experiments shown as mean +/- SD. (E) SEM images of TCs interacting with human washed platelets (WPs), thus in the absence of plasma proteins. 4T1 and B16F10 cells are highlighted, as they still show the most different platelet binding profiles (high versus low, respectively). (F) Platelet binding profiles quantification graph of (F). Data represented as mean +/-SD, from 2 to 4 independent experiments. (G) Mean of the relative number (%) of TC-bound platelets and their respective activation states, mostly discoid (resting) and filopodia (low activation). Round shapes were not observed in the absence of plasma proteins. Ring-shaped platelets of unknown function were observed in an almost undetectable frequency. Representative data from a total of 10 to 40 images per cell line from 2 to 3 independent experiments represented as mean +/- SD. (H) SEM images of TC-bound platelets at different activation states.

**Fig.S2.7.**
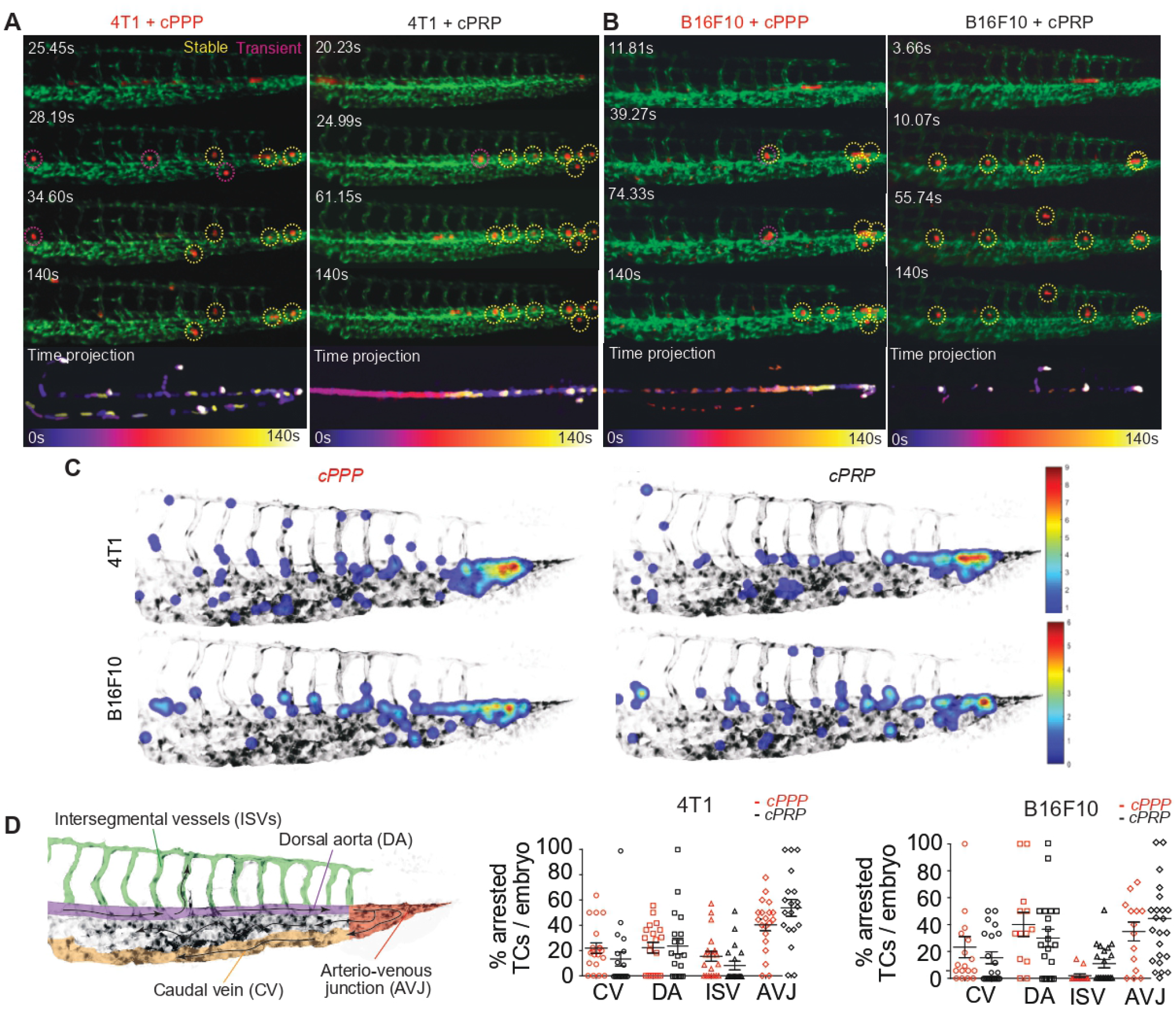
TCs interaction with platelets enhances their intravascular adhesion to the endothelial wall in a cell type-specific manner. (A) Live video microscopy time-frames and time projection of 4T1 CTCs circulation and arrest within the ZF caudal plexus in the presence (cPRP, citrated Platelet-Rich Plasma) and the absence (cPPP, citrated Plasma-Poor Plasma) of human platelets. Yellow circles highlight 4T1 cells that circulate and arrest in a sole position during the time of the assay (stable arrest). Magenta circles highlight B16F10 cells that circulate and/or arrest briefly (transient arrest). 20 embryos were analysed from 4 independent experiments. (B) Same as (A) but with B16F10. (C) Analysis by heat mapping of 4T1 and B16F10 CTCs arrest position in the presence (cPRP) or absence (cPPP) of human platelets at 5mpi. Images show TCs heatmapping analysis in Matlab as previously described5. (D) On the left, areas of the ZF circulatory system are depicted. On the right, manual counting of CTC number per ZF by vessel type (CV: caudal veins; DA: dorsal aorta; ISV: intersegmental vessels; AVJ: arterio-venous junction) in 4T1 and B16F10 models respectively. For the 4T1 model, 20 embryos were analysed from a total of 4 independent experiments. For the B16F10 model, 15 embryos from 4 independent experiments. Data are shown as mean +/- SD.

**Fig.S3.8.**
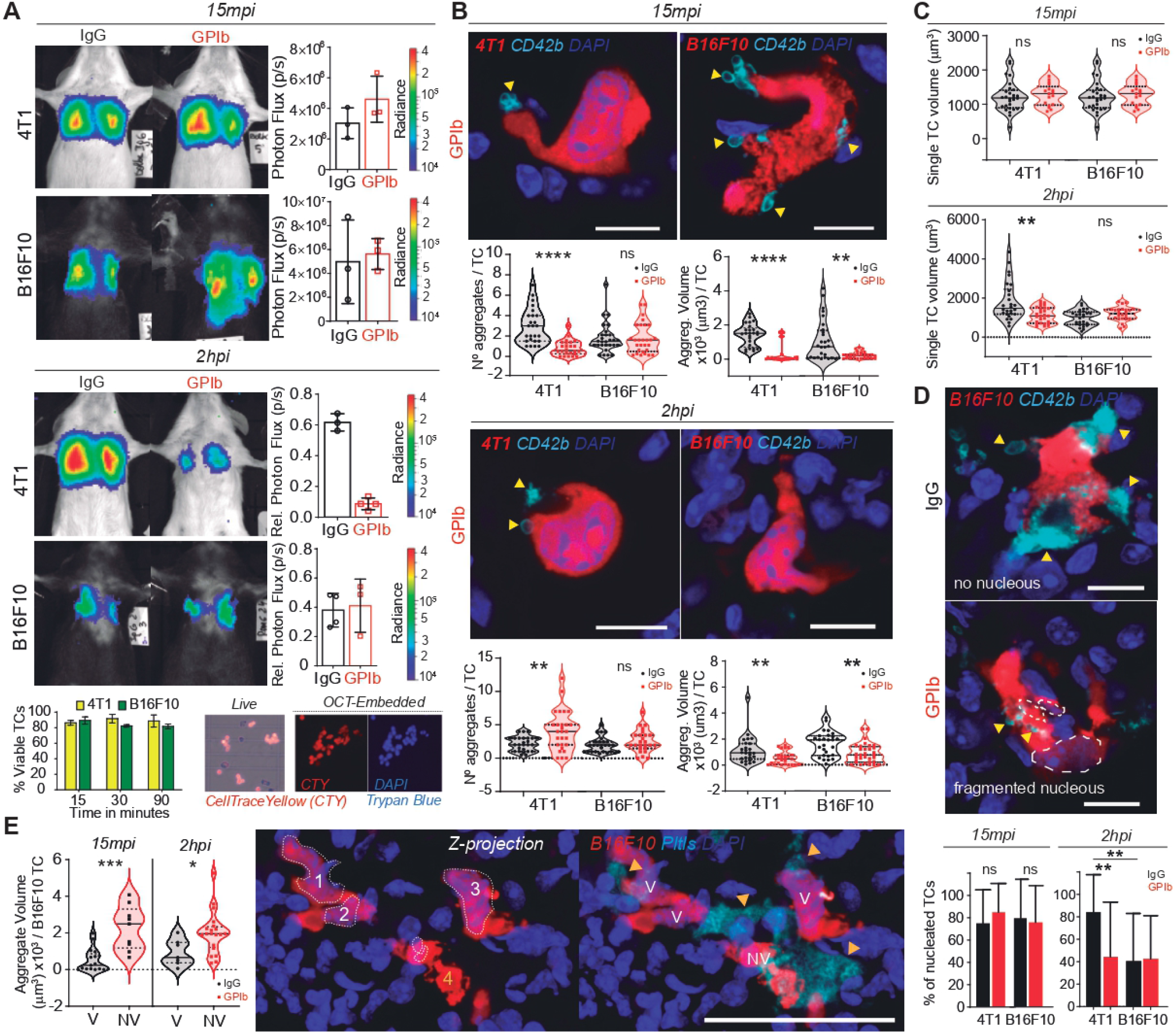
TCs’ active or passive interaction with platelets enhances their intravascular survival. (A) Representative images and time point comparisons of total lung bioluminescence in normal versus TCP mice in the 4T1 (up) and B16F10 (down) TCs models used for IF analyses of TC intravascular arrest. Two cohorts of mice were used, the first was sacrificed at 15mpi (upper panel) and the second at 2hpi (lower panel). Data are represented as mean +/- SD of 6 mice from 1 experiment. Bottom left, relative *in vitro* viabilities of 4T1 and B16F10 cells up to 90min. Bottom right, live and fluorescence imaging of and PFA-fixed / OCT-embedded 4T1 TCs labelled with CellTrace Yellow and DAPI. (B) Representative AMIRA segmentation images of single TCs arrested intravascularly in the mouse lungs after 15 minutes post-injection (upper panel) and 2hpi (lower panel). Scale: 10µm. Violin plots represent the number of aggregates per TC and the volume of aggregates per TC calculated using AMIRA segmentation of confocal images. Data are shown as mean +/- SD. (C) Violin plots depicting TCs volumes calculated in AMIRA to obtain an estimation of cell viability. Same analysis as in (B). (D) Representative images of lung-arrested non-viable B16F10 cells, as shown by the absence or fragmented nucleus. Scale: 10µm. Bar graphs (bottom) show the frequency (%) of viable B16F10 cells at 15mpi and 2hpi calculated by nuclear integrity. One experiment with n=3 different mice per group and no less than 10 single-cell images per mice. Data are shown as mean +/- SD. (E) Right: Representative confocal images of intravascular B16F10 clusters at 2hpi depicting viable cells co-opted together with non-viable cells and large platelet aggregates. Scale: 10µm. Left: Violin plots showing the volume of platelet aggregates around viable (V) and non-viable (NV) B16F10 at 15 mpi and 2phi. Same analysis as in (B).

**Fig.S4.9.**
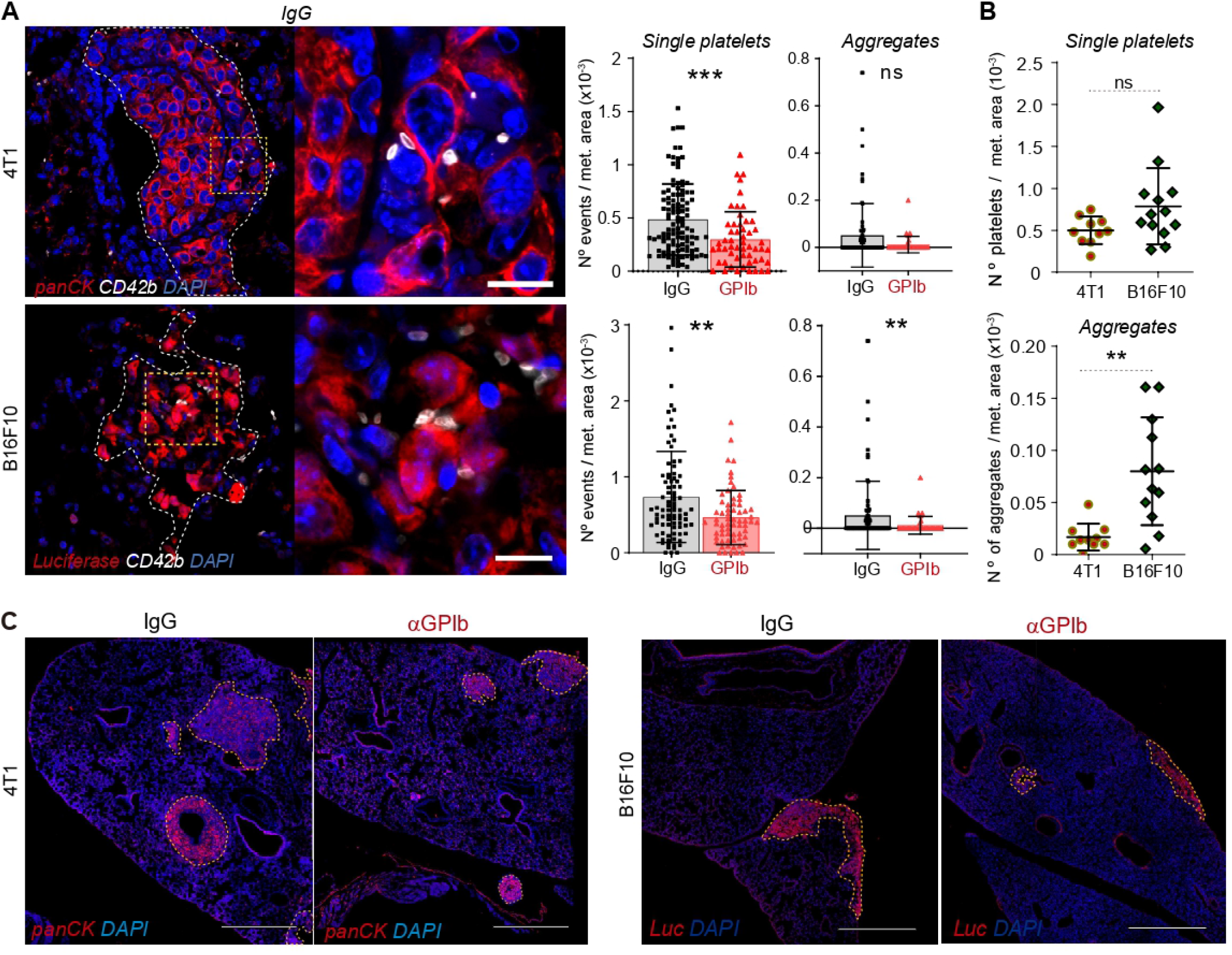
(A) Left: Representative confocal images of mouse lung metastatic foci from 4T1 (top) and B16F10 (bottom) cells at 14dpi stained for the platelet market CD42b. Scale: 50µm. Right: Bar graphs showing the number of single platelet events (< or = 3 platelets, left) and number of platelet aggregates (> 3 platelets together, right) per metastatic area, defined by α-pancytokeratin (4T1, top) or α-luciferase staining (B16F10, bottom). (B) Bar graphs comparing the number of platelets (top) and platelet aggregates (bottom) per metastatic area found in control animals of both 4T1 and B16F10 models. Data are represented as the mean +/- SD of at least 10 images per mice, from a total of 3 to 5 mice, from 3 independent experiments. (C) Low magnification confocal images stitches of lung metastases defined by pancytokeratin (4T1) or luciferase (B16F10) staining at 14dpi of control, short-term TCP mouse models injected with both 4T1 (left) and B16F10 (right) TCs. Scale bar: 500µm.

**Fig.S5.10.**
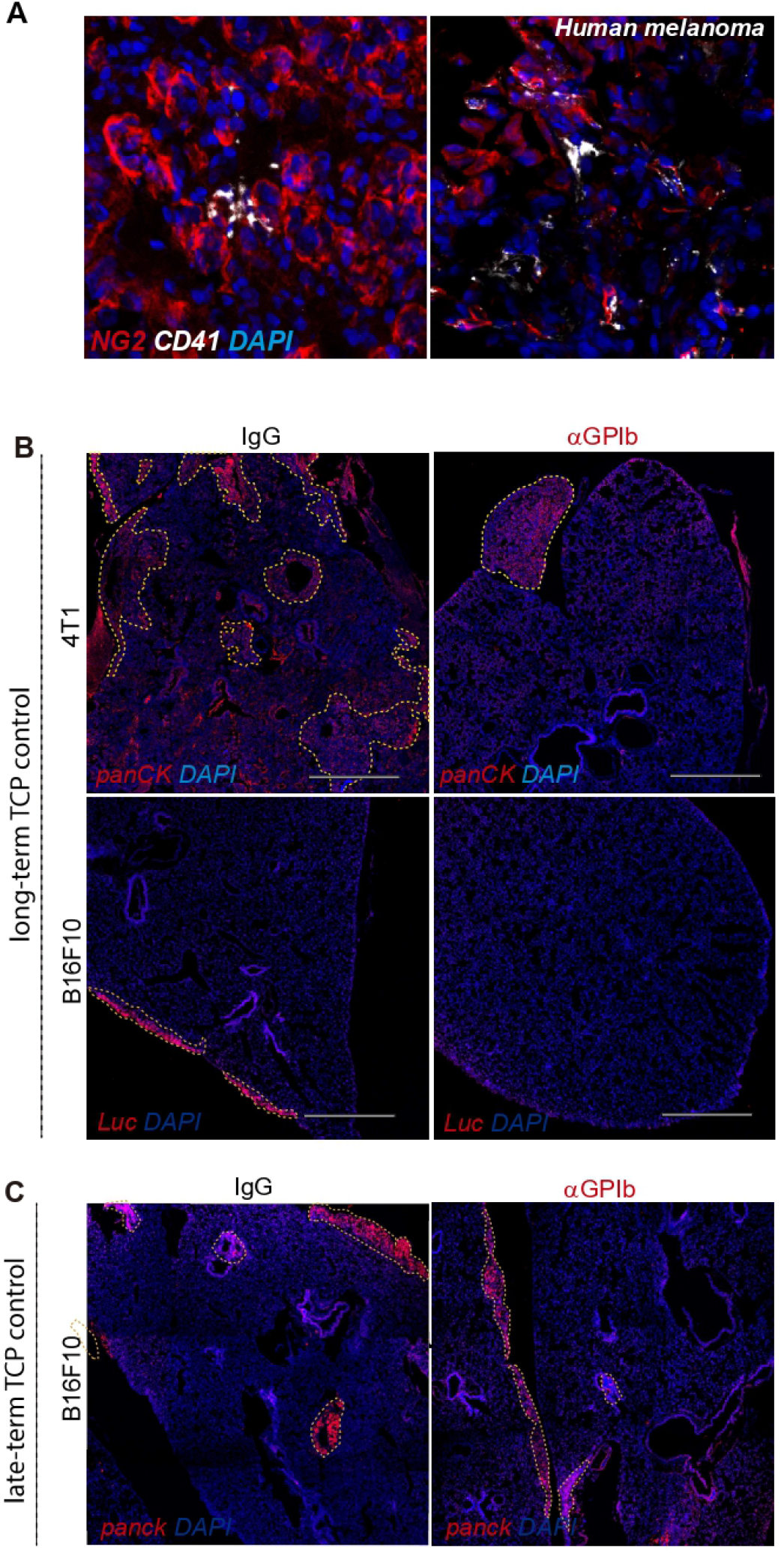
Platelets are present within lateTCP mouse and human lung metastatic foci. (A) Representative confocal images of human biopsies from melanoma metastases revealing the presence of intrametastatic platelets and platelet aggregates after staining with the platelet marker α-CD41 and melanoma marker α-NG2. (B) Low magnification confocal images of lung metastases defined by pan-cytokeratin (4T1) or luciferase (B16F10) stainings at day 14 post-injection in the long-term TCP mouse model. Scale bar: 500µm. (C) As in (B), but for late-term TCP and only for B16F10) TCs.

**Fig.S6.11.**
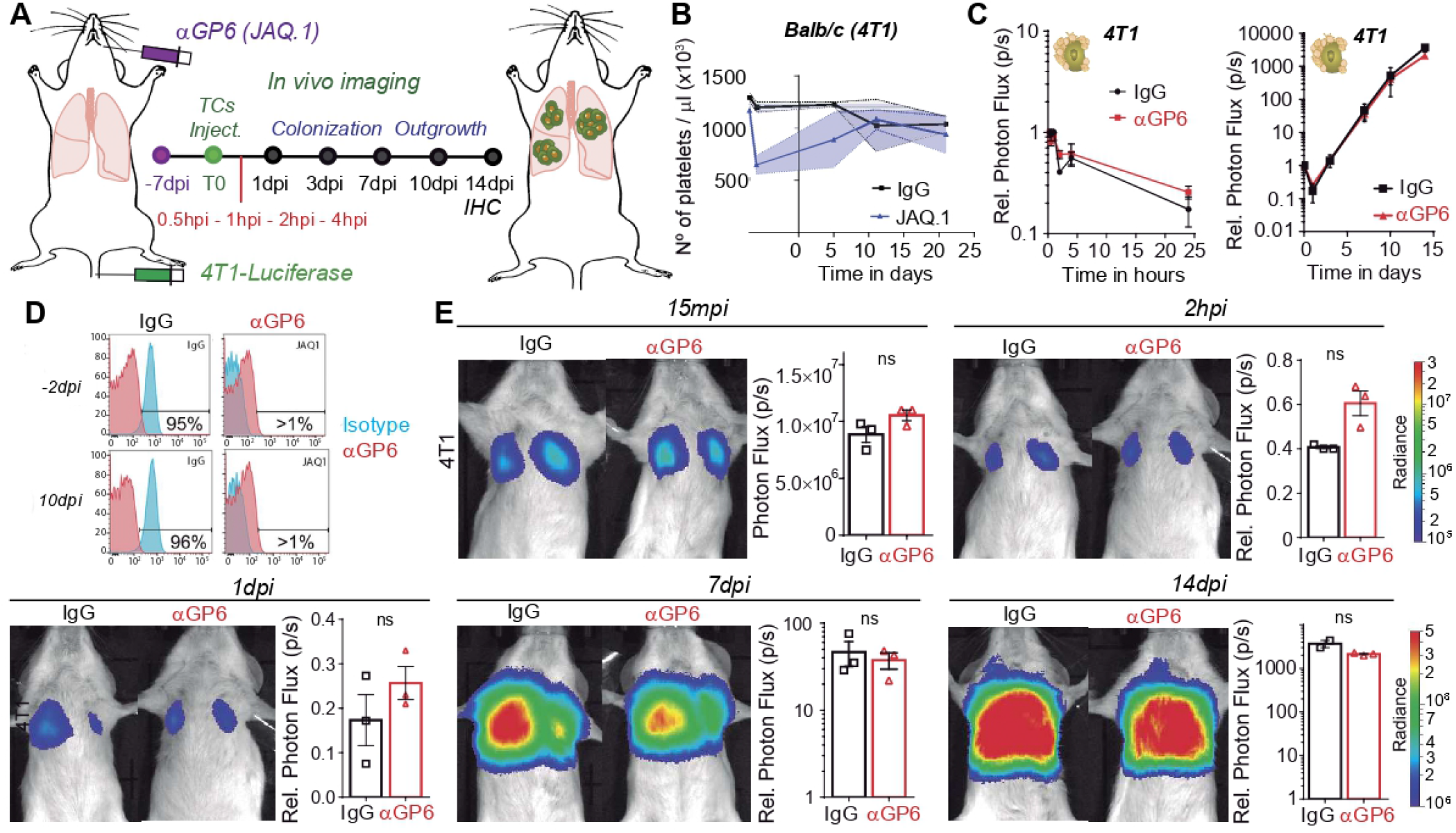
Pharmacologic blockage of the GPVI receptor does not affect lung seeding and metastatic potential of high-binding profile TCs. (A) Infographics showing the experimental setting to study the role of platelet receptor GPVI in the early and late events of the metastatic cascade (14 days) by pharmacological intervention. 4T1 TCs are injected intravenously in normal or α-GPVI-treated mice (one week before) with two I.V. shoots of the GPVI-depleting antibody JAQ.1. Lung seeding and metastatic outgrowth is followed by *in vivo* bioluminescence during 14 days. (B) Platelets counts in normal vs JAQ.1-treated animals. Despite GPVI depletion, mice show normal platelet counts after 10 days post-injection. (C) Bioluminescence kinetics of 4T1-luc TCs lung seeding (left) and outgrowth (right) in normal and α-GPVI-depleted mice. Data are presented as mean +/- SEM, 1 independent experiment of n=3. (D) Specific depletion of GPVI from platelets was assessed by flow cytometry. (E) Representative images and time point (15mpi to 14dpi) comparisons of total lung bioluminescence in normal versus GPVI-depleted animals. Data are represented as mean +/- SEM of 3 mice per group from 1 experiment.

